# Negative autoregulation mitigates collateral RNase activity of repeat-targeting CRISPR-Cas13d in mammalian cells

**DOI:** 10.1101/2021.12.20.473384

**Authors:** Chase P. Kelley, Maja C. Haerle, Eric T. Wang

**Affiliations:** Department of Molecular Genetics & Microbiology, Center for NeuroGenetics, Genetics Institute, University of Florida, Gainesville, FL; Genetics and Genomics Graduate Program, University of Florida, Gainesville, FL; Myology Institute, University of Florida, Gainesville, FL

## Abstract

Cas13 is a unique family of CRISPR endonucleases exhibiting programmable binding and cleavage of RNAs and is a strong candidate for eukaryotic RNA knockdown in the laboratory and the clinic. However, sequence-specific binding of Cas13 to the target RNA unleashes non-specific bystander RNA cleavage, or collateral activity, which may confound knockdown experiments and raises concerns for therapeutic applications. Although conserved across orthologs and robust in cell-free and bacterial environments, the extent of collateral activity in mammalian cells remains disputed. Here, we investigate Cas13d collateral activity in the context of an RNA-targeting therapy for myotonic dystrophy type 1, a disease caused by a transcribed long CTG repeat expansion. We find that when targeting CUG_n_ RNA in HeLa and other cell lines, Cas13d depletes endogenous and transgenic RNAs, interferes with critical cellular processes, and activates stress response and apoptosis pathways. We also observe collateral effects when targeting other repetitive and unique transgenic sequences, and we provide evidence for collateral activity when targeting highly expressed endogenous transcripts. To minimize collateral activity for repeat-targeting Cas13d therapeutics, we introduce <u>g</u>RNA <u>e</u>xcision for <u>n</u>egative-autoregulatory <u>o</u>ptimization (GENO), a simple strategy that leverages crRNA processing to control Cas13d expression and is easily integrated into an AAV gene therapy. We argue that thorough assessment of collateral activity is necessary when applying Cas13d in mammalian cells and that implementation of GENO illustrates the advantages of compact and universally robust regulatory systems for Cas-based gene therapies.

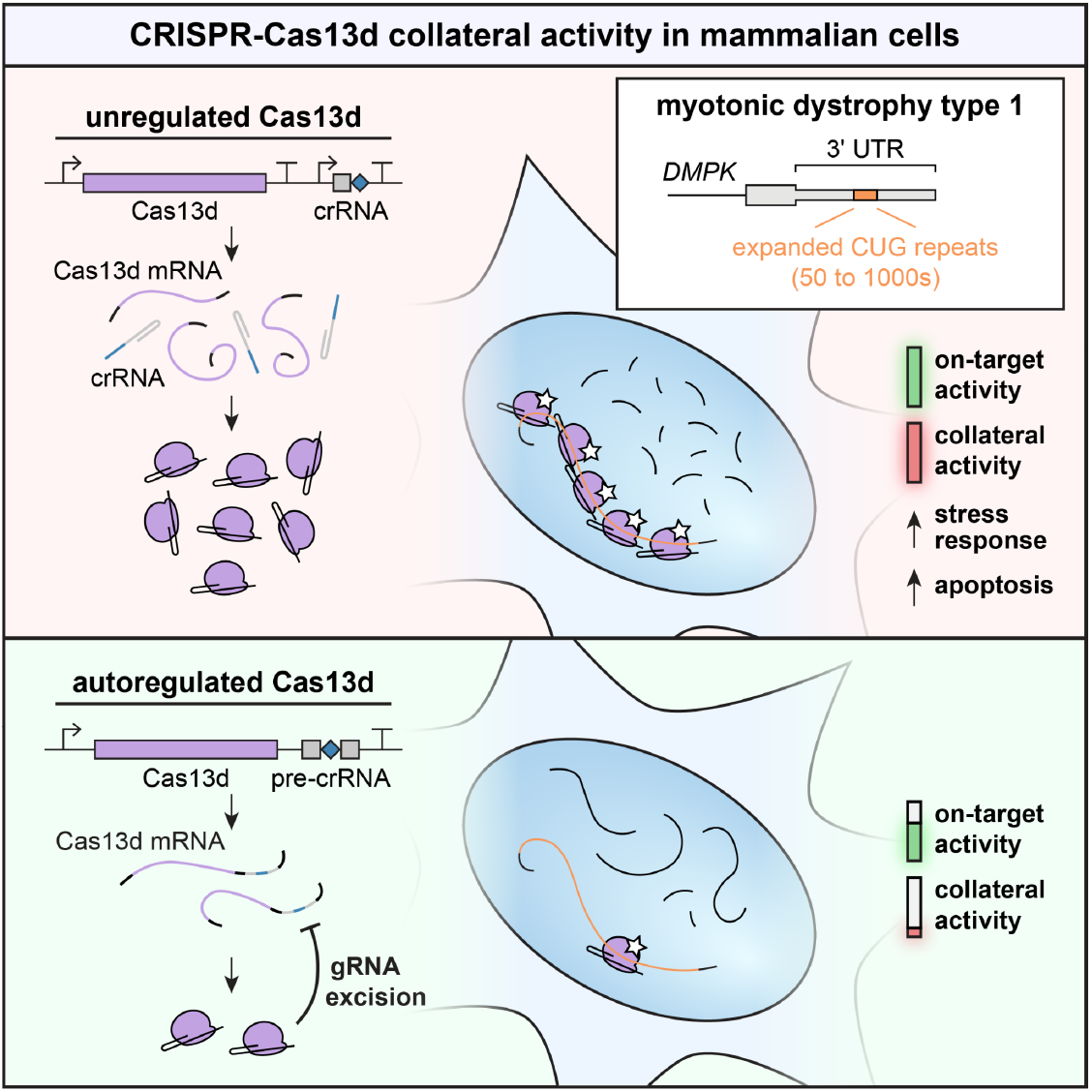

## INTRODUCTION

CRISPR-Cas13 is a recently discovered family of RNA-guided RNA endonucleases capable of sequence-specific binding and potent RNA cleavage (Abudayyeh et al., 2016). In class 2 type VI CRISPR systems, Cas13 effectors confer bacterial immunity to phage through recognition of a target RNA sequence complementary to the spacer of the crRNA, or guide RNA (gRNA) (O’Connell, 2019). Upon target binding, two higher eukaryotes and prokaryotes nucleotide (HEPN) domains undergo conformational change, forming a dual–R-X_4_-H catalytic site distal to the RNA binding cleft that exhibits potent ribonuclease (RNase) activity (Zhang et al., 2019, 2018). The programmable nature of its RNA targeting has enabled broad application of Cas13, as well as nuclease-deactivated Cas13 (dCas13) variants and fusion proteins, in eukaryotic cells to reduce expression of endogenous transcripts (Abudayyeh et al., 2017; Kushawah et al., 2020; Li et al., 2021b), introduce directed base edits (Cox et al., 2017; Kannan et al., 2021; Xu et al., 2021) and modifications (Wilson et al., 2020), visualize RNAs in live cells (Yang et al., 2019), modulate alternative splicing (Du et al., 2020; Konermann et al., 2018), and capture RNA-protein interactions (Han et al., 2020). In particular, Cas13d is a family of small orthologs that is especially suited for mammalian applications, with efficient AAV packaging for viral delivery (Konermann et al., 2018), well-studied determinants of gRNA activity (Wessels et al., 2020), and no protospacer flanking sequence (PFS) constraints (Yan et al., 2018). As a result, Cas13d is a promising candidate for a novel class of RNA-targeting therapies that avoids the potential risks of permanent genome editing or DNA binding.

A peculiar feature observed in biochemical and bacterial contexts is that, upon sequence-specific binding of the Cas13:gRNA binary complex to the target RNA, Cas13 unleashes non-specific RNase activity capable of cleaving bystander RNAs in *trans* (Abudayyeh et al., 2016; East-Seletsky et al., 2016). This phenomenon, often referred to as collateral activity, is a robust feature of all known Cas13 orthologs (O’Connell, 2019) and has been leveraged for rapid detection of nucleic acids at attomolar sensitivity (Gootenberg et al., 2017). Despite this, the extent of collateral activity of Cas13 in mammalian cells remains disputed. Many groups have observed no evidence of collateral activity in eukaryotic cells using a variety of experimental strategies (Abudayyeh et al., 2017; Huynh et al., 2020; Konermann et al., 2018; Kushawah et al., 2020), and Cas13 has been used effectively in other studies without mention of any *trans* cleavage effects (Cox et al., 2017; He et al., 2020; Li et al., 2021b; Wessels et al., 2020; Zhou et al., 2020). Yet, there is a small but growing body of evidence that collateral activity in mammalian cells substantially depletes cellular RNAs (Özcan et al., 2021; Wang et al., 2021a, 2019b; Xu et al., 2021) and that Cas13 is toxic in eukaryotes (Buchman et al., 2020). The ambiguity surrounding collateral activity calls into question the utility of Cas13 in the laboratory and presents substantial risk to safety and efficacy of Cas13-based therapeutics.

In particular, the risks of collateral activity may be magnified in development of Cas13 therapies for repeat expansion diseases, in which it can be advantageous to target the repeated sequence directly (Hu et al., 2009; Lee et al., 2012; Mulders et al., 2009). One example of this class of diseases is myotonic dystrophy type 1 (DM1), a progressive genetic disorder with multisystemic symptoms including myotonia, muscle wasting, and hypersomnolence (Ranum and Cooper, 2006). DM1 is caused by a CTG repeat expansion in the 3’ UTR of *DMPK* (Brook et al., 1992; Fu et al., 1992; Mahadevan et al., 1992), which exerts toxicity primarily through RNA gain-of-function mechanisms including sequestration of muscleblind-like (MBNL) RNA-binding proteins (Miller et al., 2000), upregulation of CELF1 (Kuyumcu-Martinez et al., 2007), and repeat-associated non-AUG (RAN) translation (Zu et al., 2011). In an ensemble of intra- and intermolecular RNA and protein interactions (Jain and Vale, 2017; Krzyzosiak et al., 2012; Querido et al., 2011), MBNL proteins and *DMPK* mRNAs containing expanded CUG repeats cluster into nuclear foci (Miller et al., 2000; Taneja et al., 1995), preventing MBNL from regulating alternative splicing and shifting isoform ratios transcriptome-wide (Otero et al., 2021; Wang et al., 2019a). Although alleles of >50 CTG motifs are associated with diagnosis of DM1, expansion lengths vary widely across the affected population and correlate strongly with severity and age of onset (Paulson, 2018). In addition, somatic instability yields wide variation in repeat length within a patient’s own cells and across tissues (Morales et al., 2012), resulting in expansions frequently reaching up to thousands of repeat units in skeletal muscle (Thornton et al., 1994) and brain (Otero et al., 2021). These effects, coupled with differential expression of *DMPK* across tissues, create a unique profile of repeat load and toxicity for each cell within each individual. Therapeutic strategies that directly target the repeat RNA sequence may provide a simple mechanism to address this complexity by titrating probability of target cleavage, and thus therapeutic potency, with repeat length at the cellular level. In support of this, we showed that binding of the CTG expansion by deactivated *Streptococcus pyogenes* Cas9 (dCas9) inhibits transcription in a repeat-length-dependent manner (Pinto et al., 2017). Cas13a has been applied successfully to degrade CUG_n_ RNA in DM1 patient-derived cells (Zhang et al., 2020); however, Cas13a orthologs are very large and pose significant challenges to AAV delivery. With potent on-target activity, lack of PFS constraints, and efficient AAV packaging (Konermann et al., 2018; Yan et al., 2018), Cas13d programmed with repeat-targeting gRNAs may be a strong candidate for developing a therapeutic platform for DM1 and other repeat expansion diseases with RNA or protein gain-of-function mechanisms. Yet, if present, collateral activity of Cas13d may be exacerbated by the many repeated and overlapping protospacers on each target RNA.

Here, we investigate the extent of collateral activity of Cas13d in mammalian cells in the context of a CUG-targeting therapy for DM1. In cell culture experiments, we find that Cas13d is effective at depleting toxic CUG_480_ RNA and ameliorating MBNL sequestration. However, in both human and mouse cell lines, we show that Cas13d collateral activity dramatically depletes orthogonal reporter transcripts when targeting CUG_480_ RNA as well as other repetitive and non-repetitive transgenic sequences. We also show evidence of collateral activity when targeting mRNAs from highly-expressed endogenous genes at unique protospacers. To combat collateral activity for repeat-targeting Cas13d therapies, we introduce <u>g</u>RNA <u>e</u>xcision for <u>n</u>egative-autoregulatory <u>o</u>ptimization (GENO), a strategy utilizing crRNA processing to minimize and stabilize Cas13d expression that can be easily implemented within AAV packaging constraints. We show that GENO substantially reduces collateral activity of Cas13d in human cells while retaining modest on-target efficacy for CUG_n_ target RNA in patient-derived DM1 myoblasts. We believe that these observations and developments help shed light on the enzymatic properties of Cas13d in mammalian cells and illuminate properties desirable in any therapeutic approach employing RNA-targeting enzymes to treat disease.

## RESULTS

### Cas13d efficiently reduces accumulation of toxic CUG_480_ RNA in nuclear foci and rescues MBNL-dependent alternative splicing

We first sought to evaluate the potential of a CRISPR-Cas13d therapeutic platform for DM1 by targeting CUG repeat RNA in a HeLa cell model and assessing the ability of Cas13d to disrupt nuclear RNA foci and rescue MBNL-mediated alternative splicing (Fig. 1A). We transfected HeLa cells simultaneously with plasmids expressing influenza hemagglutinin-(HA) tagged *Ruminococcus flavefaciens* Cas13d (RfxCas13d) with two simian virus 40 large T antigen (SV40) nuclear localization signals (NLS) and an unfused EGFP marker, a target RNA consisting of exons 11 to 15 of *DMPK* containing 480 CUG repeats (CUG_480_), and a CUG-targeting or non-targeting guide RNA (gRNA) (Fig. 1A). As Cas13d exhibits no observable protospacer flanking sequence (PFS) requirements (Konermann et al., 2018; Yan et al., 2018), we hypothesized that gRNAs targeting all three registers of the CUG repeat would initiate RNA cleavage (Fig. 1B, see Supplemental Table 1).

**Figure 1.**
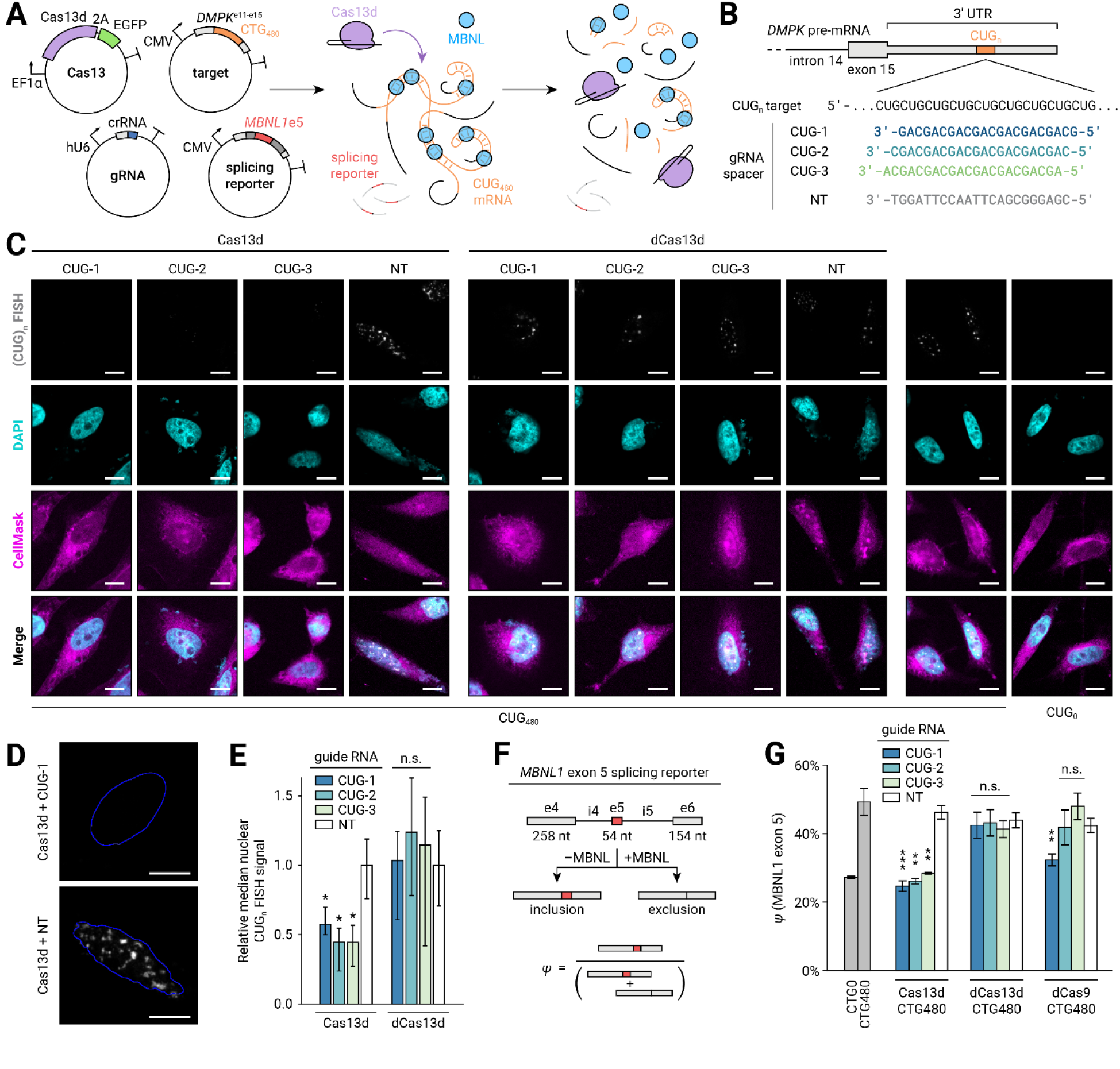
Cas13d reduces accumulation of CUG_n_ repeat RNA and rescues MBNL-dependent alternative splicing in a HeLa cell culture model of DM1. **A** Diagram of cell culture cotransfection experiments to evaluate Cas13d on-target activity and MBNL-dependent splicing. Plasmids encoding Cas13d and EGFP, gRNA, CUG_480_ target RNA, and an *MBNL1* exon 5 splicing minigene are transfected into HeLa cells and expressed before CUG_n_ RNA FISH and RT-PCR. **B** Illustration of gRNAs evaluated in this study, including three gRNAs targeting the CUG_n_ RNA and a non-targeting (NT) control. **C** Representative FISH images for CUG_n_ RNA (greyscale) in targeting and non-targeting conditions. Cells are stained with DAPI (nuclei, cyan) and CellMask (plasma membrane, magenta) for segmentation. Scale bars 10 μm. **D** Magnified images of CUG_n_ RNA FISH (greyscale) from panel C in Cas13d targeting and NT conditions. Outlines of nuclei computed by automatic thresholding are shown (blue). Scale bars 10 μm. **E** Median CUG_n_ FISH signal per nucleus, relative to NT. Error bars indicate 95% CI, estimated by bootstrapping. n>7 nuclei per condition. *p<0.05, non-overlapping CI. n.s.: not significant, p>0.05. **F** Diagram of splicing minigene assay. Sequestration of MBNL by CUG_n_ RNA increases *MBNL1* exon 5 inclusion ratio (*ψ*). Effective targeting by Cas13d aims to rescue *ψ* to unperturbed levels. **G** *MBNL1* exon 5 *ψ* for targeting and non-targeting conditions, measured by RT-PCR and capillary electrophoresis. Transcriptional inhibition by deactivated *Streptococcus pyogenes* Cas9 (dCas9) with matching spacer sequences is also shown. Error bars indicate standard deviation. n=3, all conditions. *p<0.05, **p<0.01, ***p<0.001, two-tailed Student’s *t* test. n.s.: not significant, p>0.05.

We investigated the on-target efficacy of Cas13d by performing fluorescence *in situ* hybridization (FISH) for the CUG_480_ RNA to measure the disruption of nuclear CUG RNA foci (Fig. 1C, 1D). We quantitated the mean FISH signal in each transfected nucleus across multiple widefield epifluorescence images per condition using an automated analysis pipeline (Fig. 1E). As expected, we observed a stark reduction in the intensity of nuclear foci and median nuclear FISH signal when Cas13d was paired with any of the three repeat-targeting gRNAs compared to a non-targeting gRNA (p<0.05, non-overlapping 95% confidence intervals (CI)). We did not observe significant reduction of nuclear FISH signal by nuclease-deactivated Cas13d (dCas13d) (Konermann et al., 2018) when expressed with any CUG-targeting gRNA (p>0.05, overlapping CI), indicating that target-activated RNA cleavage, rather than binding alone, is required to reduce accumulation of nuclear CUG_480_ RNA in this setting.

To evaluate the downstream impact of knockdown of CUG repeat RNA, we measured MBNL-dependent alternative splicing using a minigene consisting of *MBNL1* exons 4 through 6 with intervening introns (Fig. 1F). Inclusion of *MBNL1* exon 5 is suppressed by high concentration of available MBNL proteins in the nucleus but instead is promoted by sequestration of MBNL on CUG_n_ RNA (Gates et al., 2011). We transfected this splicing minigene along with the Cas13d, CUG_480_ target, and gRNA plasmids into HeLa cells and measured the ratio of inclusion of *MBNL1* exon 5 (*ψ*) by RT-PCR and capillary electrophoresis. We found that targeting the CUG_480_ transcript with Cas13d and any CUG-targeting gRNA significantly reduced *MBNL1* exon 5 inclusion compared to a non-targeting gRNA (Fig. 1G, p<0.01, two-tailed Student’s *t* test), suggesting that disruption of CUG_480_ RNA by Cas13d robustly reestablishes normal MBNL-dependent splicing in this model. As expected, we did not observe similar reduction of exon inclusion when gRNAs were transfected in the absence of Cas13d (Supplemental Fig. 1A), indicating that the splicing rescue observed upon Cas13d targeting is not a result of activation of endogenous RNA interference machinery.

We also did not observe any reduction of *MBNL1* exon 5 inclusion with nuclease-inactive dCas13d (Fig. 1G, p>0.05, two-tailed Student’s *t* test). To assess if dCas13d is able to successfully bind the CUG_480_ RNA, we performed FISH simultaneously with immunofluorescence (IF) using antibodies against MBNL1 and HA-tagged dCas13d (Supplemental Fig. 1B). We observed that colocalization of dCas13d with CUG_480_ RNA foci in the nucleus increased when paired with CUG-targeting gRNA rather than non-targeting gRNA (1.93 vs. 1.27 enrichment ratio, p<0.05, two-sided Mann-Whitney *U* test, Supplemental Fig. 1C), indicating that dCas13d successfully binds expanded CUG RNA in a sequence-specific manner. Intriguingly, we found that enrichment of MBNL1 in foci appeared only partially reduced by CUG-targeting dCas13d (1.97 vs. 2.90 enrichment ratio, p=0.057, two-sided Mann-Whitney *U* test, Supplemental Fig. 1C). These results suggest that although dCas13d may compete with MBNL1 at binding sites within CUG RNA foci, partial displacement is insufficient to restore MBNL1-mediated splicing in this model system.

We have previously shown that nuclease-deactivated *Streptococcus pyogenes* Cas9 (dCas9) paired with a CTG-targeting gRNA efficiently blocks transcription of CUG_n_ RNA (Pinto et al., 2017). We cloned the dCas9 gene into an expression context matching that of Cas13d, and we transfected this dCas9 plasmid into HeLa cells along with CUG_480_ target, the *MBNL1* exon 5 minigene, and CTG-targeting or non-targeting gRNA with spacers matching our Cas13d gRNAs. As expected, dCas9 rescued splicing of *MBNL1* exon 5 only when paired with the CUG-1 gRNA (Fig. 1G, p<0.01, two-tailed Student’s *t* test). This gRNA aligns to the register of the CTG_480_ repeat with a CAG protospacer adjacent motif (PAM), which is acceptable for binding of dCas9 (Hsu et al., 2013; Jiang et al., 2013). gRNAs targeting registers with AGC or GCA PAMs (CUG-2 and CUG-3, respectively) did not rescue MBNL alternative splicing (Fig. 1G, p>0.05, two-tailed Student’s *t* test). In contrast, all three registers supported rescue of splicing by Cas13d, suggesting that the lack of PFS requirements for binding and activation of Cas13d expands the available sequence targeting space for CRISPR therapies.

### CUG-targeted Cas13d inhibits EGFP fluorescence

Although Cas13d was effective at reducing CUG_480_ RNA accumulation and restoring MBNL-mediated splicing, we noticed that we were unable to visualize expression of the EGFP marker after transfection of Cas13d, CUG-targeting gRNA, and CUG_480_ target plasmids, but EGFP expression was apparent when either dCas13d or non-targeting gRNA were substituted (Supplemental Fig. 2A). To quantitate this phenomenon, we transfected CUG_480_ target plasmid into HeLa cells along with either Cas13d or dCas13d and with either CUG-targeting or non-targeting gRNA, and we measured EGFP fluorescence 20 hr after transfection. We found that EGFP expression was reduced by >96% upon transfection with Cas13d and any CUG-targeting gRNA compared to the non-targeting control (Supplemental Fig. 2B, p<0.05, two-tailed Student’s *t* test). The same effect was not observed for dCas13d (Supplemental Fig. 2B), indicating that loss of EGFP fluorescence results from Cas13d RNase activity.

To ensure that this reduction of EGFP fluorescence does not merely reflect differences in cell survival, we performed a resazurin cell viability assay 20 hr and 44 hr after transfection. After 20 hr, we observed only a slight reduction in cell viability when Cas13d was transfected with CUG-1 or CUG-2 gRNA compared to non-targeting gRNA (Supplemental Fig. 2C, 8.0% reduction for CUG-1 and 4.8% for CUG-2, p<0.01, two-tailed Student’s *t* test), indicating that the observed loss of EGFP signal after 20 hr is not explained by death of transfected cells. However, a larger reduction in cell viability was observed 44 hr after transfection for all CUG-targeting gRNAs (Supplemental Fig. 2C, >16% reduction for all gRNAs, p<0.001, two-tailed Student’s *t* test), suggesting that persistent targeting of CUG_n_ RNA by Cas13d is cytotoxic.

### CUG-targeted Cas13d upregulates stress response and apoptosis signaling pathways

To further investigate these observations, we performed RNA sequencing (RNA-seq) to profile biological pathways disrupted by CUG-targeting Cas13d and to compare these transcriptomic changes to effects of other repeat-targeted technologies (Supplemental Fig. 3). We transfected HeLa cells with Cas13d and with or without CUG-1 gRNA and sequenced RNA collected 68 hr after transfection (Supplemental Fig. 3A). For comparison, we also tested dCas9 with and without CUG-1 gRNA as well as sequence-matched CUG-targeting and non-targeting short hairpin RNAs (shRNAs), which utilize the endogenous RNA interference machinery for knockdown (Paddison et al., 2002) and are frequently used as comparators for Cas13 off-targeting (Abudayyeh et al., 2017). Transcripts from 57 genes in the human reference genome (hg19) contain CUG_n_ repeats that extend beyond the length of the Cas13d spacer (Supplemental Table 2) and therefore are likely off-targets of CUG-targeting approaches. We omitted the CUG_480_ target plasmid in this experiment to enrich for these off-target events. We observed consistently strong correlations of transcripts per million (TPM) estimates between RNA-seq libraries across all conditions (Supplemental Fig. 3B, minimum Pearson *r* of log(TPM) = 0.92, median = 0.98), suggesting low variance introduced during library preparation.

We found that CUG-targeting Cas13d disrupted fewer off-target genes than shRNA (116 vs. 443 genes differentially expressed (DE) between targeting and non-targeting conditions, Benjamini-Hochberg false discovery rate (FDR) q<0.05) but many more genes than dCas9 (3 DE genes, FDR q<0.05) (Supplemental Fig. 3C). 69% of genes disrupted by CUG-targeting shRNA were downregulated (304 vs. 139 genes), consistent with the notion that RISC-mediated RNA cleavage is the dominant mechanism underlying differential expression. In contrast, most of the genes perturbed by Cas13d were upregulated (83 vs. 33 genes), suggesting that a mechanism other than *cis* RNA cleavage is responsible for most DE genes.

Of the genes disrupted by dCas9, two were downregulated (*CALM1* and *ITGA5,* 30% reduction) and one was upregulated (*ANKMY2,* 70% increase). The 5’ UTR of *CALM1* contains a perfectly matching protospacer with a CGG PAM, and *ITGA5* exon 1 contains a protospacer with a single mismatch and CGG PAM. Both prospective targets fall <150 bp from the transcription start site (TSS), suggesting steric inhibition of transcription elongation. A perfect protospacer with a TGG PAM is located 385 bp upstream of the TSS of *ANKMY2* and falls within an H3K27ac-enriched active enhancer, suggesting that dCas9 may interact with *cis* or *trans* factors within this enhancer to increase transcription. In all three cases, prospective dCas9 target sites found within or near off-target gene loci strongly suggest that differential expression is driven by sequence-specific interactions.

To determine if the differences in off-target profiles could reflect differences in the strength of knockdown of RNAs containing CUG repeats, we binned transcripts by the length of their longest CUG repeat in the hg19 reference genome and calculated the median fold change between targeting and non-targeting conditions. We found a strong relationship between off-target knockdown by shRNA and CUG repeat length for genes with repeats shorter than the shRNA targeting sequence (Supplemental Fig. 3D), and we observed an average of 35% knockdown of transcripts with CUG repeats >22 nt (Supplemental Fig. 3E). In contrast, both Cas13d and dCas9 exhibited substantially reduced knockdown of off-target RNAs containing short CUG repeats, with very similar profiles between the two CRISPR effectors (Supplemental Fig. 3D). In fact, the median knockdown of transcripts with CUG repeats >22 nt was slightly weaker for Cas13d than dCas9 (6.4% vs. 8.9%, Supplemental Fig. 3E). This suggests that knockdown of endogenous genes containing CUG repeats does not fully explain the extensive off-target profile of CUG-targeted Cas13d.

We performed gene ontology (GO) analysis using PANTHER (Mi et al., 2019) to investigate the biological pathways enriched in these differentially expressed genes. We found that the majority of biological processes perturbed by Cas13d were involved in either stress response (48%) or apoptosis (16%) signaling pathways (Supplemental Fig. 3F, enrichment >5, FDR q<0.05). In contrast, we observed reduced enrichment of stress response pathways (18% of enriched processes) and lack of enrichment of apoptosis signaling when cells were treated with CUG-targeting shRNAs (Supplemental Fig. 3G). These results are consistent with the notion that Cas13d activation by CUG repeats leads to toxicity and cell death beyond what is expected from *cis* cleavage of off-target RNAs alone.

### CUG-targeted Cas13d reduces expression of orthogonal mCherry reporter in mammalian cells

We hypothesized that inhibition of EGFP expression and induction of stress response genes may result from global depletion of cellular RNAs by Cas13d collateral RNase activity upon activation by CUG_n_ RNA. To assay for collateral activity, we cotransfected a plasmid constitutively expressing mCherry as an orthogonal reporter gene into HeLa cells along with Cas13d, gRNA, and CUG_480_ target plasmids, and we measured mCherry fluorescence in response to Cas13d targeting of CUG_480_ RNA (Fig. 2A). We reasoned that if Cas13d collateral activity is extensive, we should expect to observe a general reduction of cellular transcripts, including transgenic mCherry mRNA, which would reduce subsequent translation and fluorescence (Fig. 2B). We measured mCherry fluorescence 20 hr after transfection to minimize the impact of cell viability on bulk fluorescence measurements.

**Figure 2.**
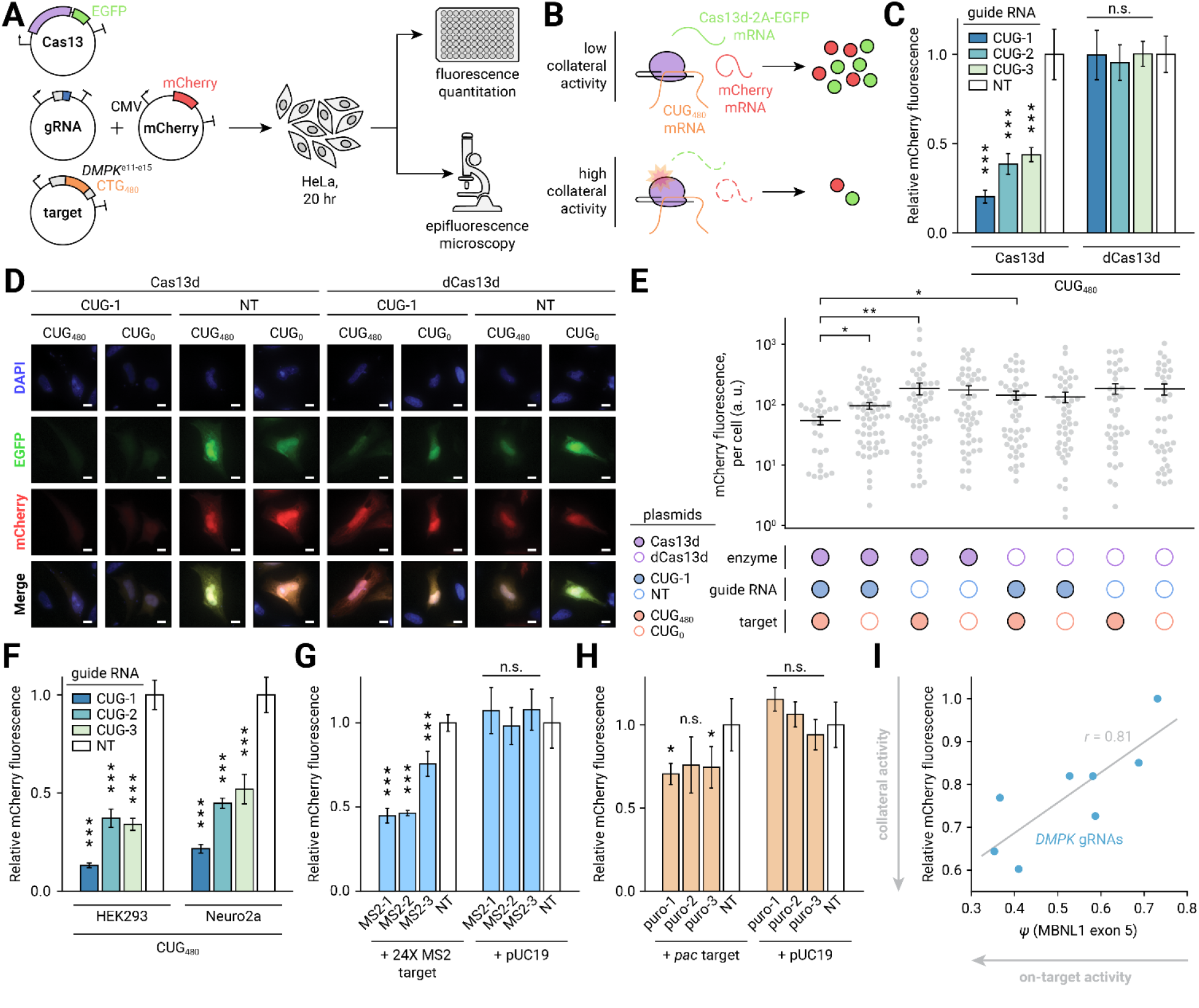
Activation of Cas13d reduces expression of orthogonal reporter genes in eukaryotic cells. **A** Diagram of fluorescence assays to detect collateral activity of Cas13d. HeLa cells are transfected with plasmids encoding Cas13d and EGFP, gRNA, target RNA, and a constitutively expressed orthogonal mCherry reporter. After 20 hr, mCherry fluorescence is measured by plate reader or epifluorescence microscopy. **B** Interpretation of fluorescence assays. If collateral activity is weak, expression levels of EGFP and mCherry reporter genes are unaffected by Cas13d activation. If collateral activity is strong, both mCherry and EGFP gene products are depleted in *trans* upon activation. **C** Bulk quantitation of mCherry fluorescence by plate reader in CUG-targeting and non-targeting (NT) conditions, relative to NT. Error bars indicate standard deviation. n=5, all conditions. ***p<0.001, two-tailed Student’s *t* test. n.s.: not significant, p>0.05. **D** Representative images from epifluorescence microscopy of mCherry and EGFP reporter genes in CUG-targeting and non-targeting conditions. Nuclei are stained with DAPI. Scale bars 10 μm. **E** Quantitation of mCherry fluorescence by epifluorescence microscopy. Measurements from individual cells are shown (grey dots) along with the mean (black line). Error bars indicate SEM. *p<0.05, **p<0.01, two-sided Mann-Whitney *U* test. **F** Bulk mCherry measurement when targeting CUG_480_ RNA with Cas13d in HEK293 and Neuro2a cell lines. ***p<0.001, two-tailed Student’s *t* test. **G** Bulk mCherry measurement when targeting a transgenic transcript containing 24X MS2 hairpin sequences with Cas13d using MS2-targeting gRNAs. A control plasmid (pUC19) that does not express the MS2 target is also included. ***p<0.001, two-tailed Student’s *t* test. n.s.: not significant, p>0.05. **H** Bulk mCherry measurement when targeting a transgenic transcript encoding puromycin acetyltransferase (*pac*) with Cas13d at unique sites using *pac*-targeting gRNAs. *p<0.05, two-tailed Student’s *t* test. n.s.: not significant, p>0.05. **I** Relationship between on-target efficacy (rescue of MBNL1 exon 5 *ψ*) and mCherry inhibition for gRNAs targeting unique *DMPK* sequences present in the CUG_480_ transcript. Grey line indicates ordinary least squares linear regression (Pearson *r* = 0.81).

We observed a >56% reduction in mCherry fluorescence in cells transfected with Cas13d and any CUG-targeting gRNA relative to non-targeting gRNA (Fig. 2C, p<0.001, two-tailed Student’s *t* test). We did not observe any reduction in mCherry fluorescence with dCas13d paired with any gRNA, indicating that this effect results from Cas13d RNase activity. To assay fluorescence at the single-cell level, we performed widefield fluorescence microscopy and quantitated mCherry intensity for each cell (Fig. 2D). On average, we observed a >61% decrease in mCherry fluorescence when cells were transfected with Cas13d, CUG-1 gRNA, and CUG_480_ target compared to conditions when either dCas13d or NT gRNA were substituted and a 43% decrease compared to a condition where the target was substituted for CUG_0_ (Fig. 2E, p<0.05, one-sided Mann-Whitney *U* test). We also observed a smaller reduction in mCherry intensity (44%) between CUG-targeting and non-targeting conditions in the absence of CUG_480_ target (p<0.05, one-sided Mann-Whitney *U* test), likely due to activation of Cas13d collateral activity by endogenous transcripts. Overall, these results provide strong evidence of abundant Cas13d collateral activity in HeLa cells when targeting CUG repeat RNA.

Other researchers have observed that the extent of collateral activity of Cas13a is dependent on cell type, finding that collateral activity was apparent in human glioma cells but not detectable in the non-cancerous HEK293 line (Wang et al., 2019b). To determine if collateral activity of Cas13d is specific to HeLa cells, we performed the same experiment in HEK293 and the mouse neuroblastoma Neuro2a line. We observed similar levels of mCherry inhibition across all cell lines (Fig. 2F, p<0.001, two-tailed Student’s *t* test), suggesting that collateral activity of Cas13d in mammalian cells is a general phenomenon.

### Cas13d reduces mCherry expression in a target-dependent manner

To determine whether collateral RNase activity is specific to targeting CUG repeats, we designed three gRNAs complementary to the MS2 hairpins from a plasmid expressing 24 MS2 repeats in tandem (Supplemental Table 1). We transfected these gRNAs into HeLa cells along with Cas13d and either the 24X MS2 target plasmid or a control plasmid without MS2 hairpins (pUC19). We observed significant reduction of mCherry fluorescence with all three MS2-targeting gRNAs when the 24X MS2 target was present (Fig. 2G, average 44% reduction, p<0.001, two-tailed Student’s *t* test), but not with the control target (p>0.05, two-tailed Student’s *t* test). This experiment illustrates that collateral activity is not specific to CUG_n_ RNA targets and is only detected when suitable protospacers are present in the transcriptome.

Due to their repetitive nature, it is plausible that repeat expansion RNAs would greatly increase collateral activity by binding many Cas13d molecules simultaneously. We sought to determine whether collateral activity is also detectable at unique protospacer sequences. We designed three gRNAs that target unique sequences within the puromycin acetyltransferase (*pac*) gene (Supplemental Table 1), and we transfected these gRNAs into HeLa along with Cas13d and either a *pac*-expressing target plasmid or a control plasmid (pUC19). We detected a significant decrease in mCherry signal for two of the three gRNAs only when co-transfected with the *pac* target plasmid (Fig. 2H, average 28% reduction, p<0.05, two-tailed Student’s *t* test) and not with the control target (p>0.05, two-tailed Student’s *t* test). These data suggest that although collateral activity may be weaker at unique protospacers than repetitive targets, it may still have a deleterious impact on cellular RNAs and should be thoroughly evaluated prior to use of Cas13d for RNA knockdown in mammalian cells.

### Rescue of MBNL splicing by *DMPK*-targeted Cas13d correlates with mCherry inhibition

The above observations support the notion that collateral activity is a general property of the Cas13d enzyme, the rate of which is likely a function of many variables, including biochemical and biological context, concentration of target protospacers, and affinity between the Cas13d:gRNA binary complex and the target RNA. As *cis* RNase activity of the target RNA by Cas13d after binding has been shown to be largely independent of flanking sequences (Yan et al., 2018) (apart from a preference for uracil bases (Konermann et al., 2018)), we reasoned that both on-target and collateral RNase activities may be driven primarily by the apparent intracellular *K_D_* of the Cas13d:gRNA:target ternary complex and thus may be positively correlated for a given target RNA. To test this hypothesis, we designed eight gRNAs that target unique protospacers within the 3’ UTR of *DMPK* (all of which are present in the CUG_480_ target RNA, see Supplemental Table 1), and we co-transfected these gRNAs into HeLa cells with Cas13d, CUG_480_ target, the *MBNL1* exon 5 splicing minigene, and mCherry plasmids. To probe on-target activity, we measured the ratio of inclusion of *MBNL1* exon 5, and we measured bulk mCherry fluorescence to assay collateral activity. We found a positive correlation between MBNL splicing rescue and mCherry fluorescence (Fig. 2I, Pearson *r* = 0.81), suggesting that on-target and collateral activity of Cas13d are strongly linked.

### Collateral RNase activity of Cas13d is observed when targeting endogenous genes

Our previous observations of collateral activity were made while targeting overexpressed transgenes with Cas13d in a transient transfection format. We reasoned that the extent of collateral activity, as a biochemical process driven by the Cas13d ternary complex, would depend stoichiometrically on both Cas13d protein and target RNA concentrations. We therefore desired to assess whether collateral activity may still be detectable and/or toxic when targeting endogenous genes with genomically encoded Cas13d. We designed gRNAs against six endogenous genes across a wide range of expression levels in HeLa cells (*LDHA, CD63, CD81, LGMN, SYBU, EPOR;* see Supplemental Table 3). In a comprehensive high-throughput CRISPR screening study (Hart et al., 2015), none of these genes were classified as core fitness genes, and all had low confidence scores for essentiality in HeLa cells (Bayes factor < −10), making them strong candidates for assaying the detrimental effects of collateral activity.

In an effort to generate a cell line constitutively expressing Cas13d, we treated HeLa cells with lentivirus packaged with the Cas13d-T2A-EGFP gene under an EF1α promoter and selected GFP-positive single cells by flow cytometry, yet we were unable to identify any clonal lines without substantial truncations of Cas13d after expansion (Supplemental Fig. 4A). Instead, we generated a HeLa cell line with chemically inducible Cas13d by integrating a piggyBac transposon (Cadiñanos and Bradley, 2007) expressing Cas13d-T2A-EGFP under a tetracycline-inducible promoter (Fig. 3A). As a reporter for collateral activity, we also integrated a second cassette constitutively expressing mCherry, and we expanded a clonal line (HeLa-tet:Cas13d-mCherry) and validated the expression of full-length Cas13d by Western blot and EGFP and mCherry by fluorescence microscopy after incubation with 2 μM doxycycline for 44 hr (Supplemental Fig. 4B, 4C).

**Figure 3.**
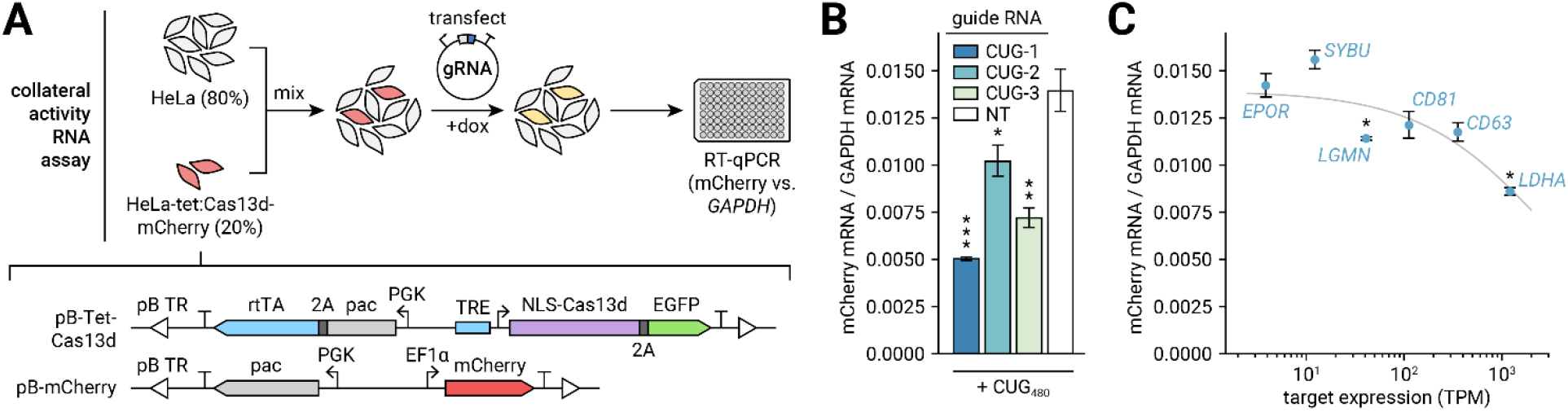
Targeting endogenous mRNAs with Cas13d activates collateral RNAse activity. **A** Description of the collateral activity RNA assay. The HeLa-tet:Cas13d-mCherry cell line expresses mCherry constitutively and Cas13d under a doxycycline-inducible promoter. As collateral activity likely depletes transgenic reporter and endogenous RNAs, non-transgenic HeLa cells are spiked in at a 4:1 ratio prior to transfection of gRNA plasmid and induction of Cas13d expression by doxycycline. After 44 hr, mCherry and control (*GAPDH*) mRNA expression is measured by RT-qPCR. **B** Assay validation using CUG-targeting gRNAs and transfected CUG_480_ target plasmid. n=3 biological replicates per condition, averaged across two technical qPCR replicates. Error bars indicate SEM of Δ*C_q_* propagated to the plotted ratio. *p<0.05, **p<0.01, ***p<0.001, one-tailed Student’s *t* test of Δ*C_q_*. **C** Ratio of mCherry to *GAPDH* mRNA expression observed when targeting endogenous genes. Average target expression was calculated from RNA-seq of untransfected HeLa cells included in our experiment described in Supplemental Fig. 3. Error bars indicate SEM of Δ*C_q_* propagated to the plotted ratio. Grey line depicts a logistic function fit to the data using non-linear least squares regression. *p<0.05, one-tailed Student’s *t* test of Δ*C_q_*.

To measure collateral RNase activity at the RNA level, we performed an RT-qPCR experiment to compare abundance of mCherry reporter and control gene (*GAPDH*) transcripts (Fig. 3A). Since we expected collateral activity to degrade RNAs nonspecifically in the transgenic cell line (including both mCherry and *GAPDH* transcripts), we mixed HeLa-tet:Cas13d-mCherry cells with nontransgenic HeLa cells at a 1:4 ratio. We transfected this co-culture with gRNA-encoding plasmids, and we induced Cas13d expression with 2 μM doxycycline for 44 hr prior to RNA extraction. As the nontransgenic HeLa cells did not express Cas13d or mCherry, they provided a stable baseline level of *GAPDH* transcripts without influencing mCherry transcript abundance, enabling measurement of collateral activity by RT-qPCR. To validate the method, we transfected CUG-targeting and non-targeting gRNAs along with CUG_480_ target, and we observed reduction of mCherry RNA abundance with all CUG-targeting gRNAs comparable to our fluorescence assays (Fig. 3B, average 46% reduction vs. NT gRNA, p<0.05, one-tailed Student’s *t* test of Δ*C_q_).*

Using this approach, we observed a statistically significant reduction of mCherry RNA when targeting two of the six endogenous genes in our panel (*LDHA* and *LGMN;* Fig. 3C, p<0.05, one-tailed Student’s *t* test of Δ*C_q_*). We also noticed an overall negative correlation between target expression level and mCherry RNA abundance (Pearson *r* = −0.87 between [mCherry] and log(target TPM)). These data suggest that although collateral activity may be strong when targeting highly expressed genes (eg. 34% depletion for *LDHA*), its effects may be difficult to detect for weakly expressed targets. From this analysis, we argue that although collateral activity of Cas13d is most problematic when targeting highly expressed and/or repetitive RNAs, it is important to screen for its effects with any target to ensure that nonspecific RNA depletion does not confound interpretation of experiments or cause unintended cellular toxicity.

To investigate whether collateral activity activated by endogenous genes is sufficient to produce toxicity, we performed a resazurin cell viability assay on HeLa-tet:Cas13d-mCherry cells transfected with gRNA plasmids and induced with 2 μM doxycycline for 44 hr. We observed statistically significant reductions in cell viability for the three most highly expressed targets (*LDHA, CD63, CD81*; Supplemental Fig. 4D, p<0.05, two-tailed Student’s *t* test) and a clear trend of increasing cell mortality with target expression level. We found that cell viability did not correlate with depletion of gRNAs targeting these genes in a high-throughput CRISPR screen in HeLa cells (Hart et al., 2015) (Supplemental Fig. 4E, p>0.05, beta distribution c.d.f.), providing further evidence that the observed toxicity was caused by collateral activity of Cas13d rather than on-target knockdown effects. It is important to note that cell viability may explain part of the observed reduction of mCherry RNA abundance (Fig. 3C), yet both measurements likely represent different manifestations of the same collateral activity phenomenon. Overall, these experiments suggest that collateral activity of Cas13d is strongest at repetitive and/or abundant targets due to stoichiometric activation of a large number of Cas13d:gRNA complexes.

### Negative autoregulation by gRNA excision reduces expression of Cas13d

We reasoned that the presence of many overlapping protospacers on a repetitive target RNA may exacerbate collateral activity by binding many copies of Cas13d:gRNA complex simultaneously (Fig. 4A). Therefore, we hypothesized that reduction of Cas13d expression may reduce collateral activity while maintaining significant on-target cleavage of the CUG_n_ RNA. We developed a negative autoregulation strategy, termed gRNA excision for negative-autoregulatory optimization (GENO), to sharply reduce and control Cas13d expression by leveraging its crRNA processing activity for self-knockdown (Fig. 4B). In GENO, a pre-crRNA containing the spacer flanked by two direct repeats is placed within a UTR of the Cas13d mRNA. After transcription and translation, Cas13d excises and processes the gRNA to form the binary complex, cleaving its mRNA in the process and leading to degradation and prevention of further translation.

**Figure 4.**
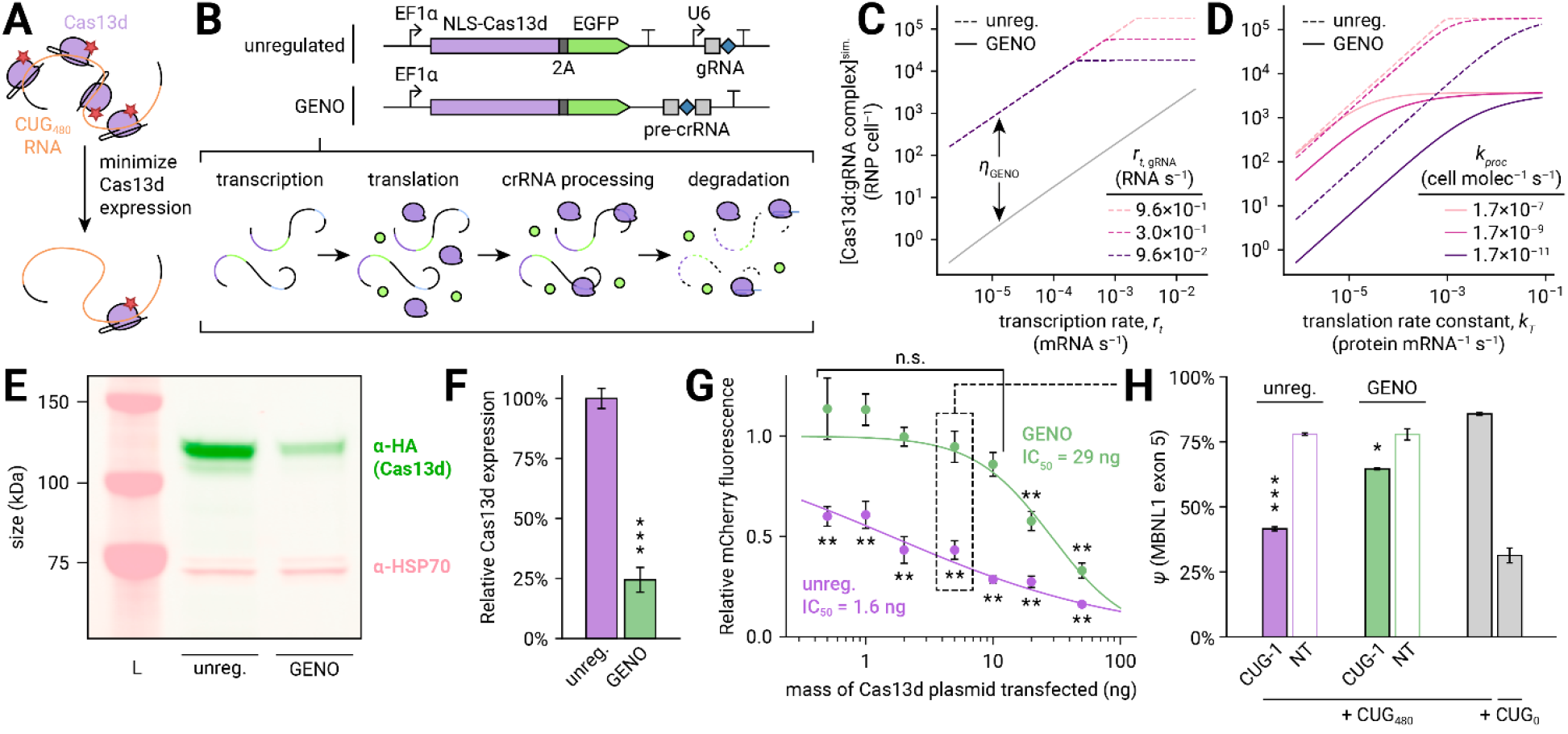
Negative autoregulation by gRNA excision reduces Cas13d collateral activity and retains on-target activity at CUG_n_ target RNA. **A** Diagram describing rationale for negative autoregulation of Cas13d. Overexpression likely produces many copies of Cas13d:gRNA binary complex for each CUG_n_ target RNA, activating many molecules of Cas13d for *trans* cleavage. Minimizing Cas13d expression may reduce collateral activity dramatically with minimal loss of on-target activity. **B** Description of gRNA excision for negative-autoregulatory optimization (GENO) approach to control Cas13d expression. In GENO, the pre-crRNA is located within a UTR of the Cas13d transcript, leading to cleavage and degradation of Cas13d mRNA during crRNA processing. **C** Simulation of steady-state concentration of Cas13d:gRNA binary complex in autoregulated (solid line) and unregulated (dashed lines) conditions as a function of Cas13d transcription rate (*r_t_*, horizontal axis) and gRNA transcription rate in the unregulated design (*r*_*t*,gRNA_, isolines). The autoregulation efficiency *η*_GENO_ is also annotated (see Supplemental Note). Translation rate constant *k_T_* = 8.3 × 10^-3^ protein mRNA^-1^ s^-1^, crRNA processing rate constant *k_proc_* =1.7 × 10^-7^ cell molec^-1^ s^-1^. **D** Simulation of Cas13d:gRNA complex concentration as a function of translation rate constant (*k_T_*, horizontal axis) and crRNA processing rate constant (*k_proc_,* isolines). Cas13d transcription rate *r_t_* = 0.02 mRNA s^-1^, gRNA transcription rate in unregulated design *r*_*t*,gRNA_ = 0.96 RNA s^-1^. **E** Fluorescent Western blot comparing Cas13d protein expression from unregulated and GENO plasmids transfected into HeLa cells. Cas13d is visualized with α-HA antibody (green), and HSP70 is stained as a loading control (pink). Protein ladder is shown (L). **F** Quantification of Cas13d Western blot. Error bars indicate standard deviation. n=3 transfections per condition. ***p<0.001, two-tailed Student’s *t* test. **G** mCherry fluorescence 20 hr after transfection of HeLa cells with unregulated (purple) and GENO (green) Cas13d and CUG-1 gRNA plasmids, CUG_480_ target, mCherry, and pUC19 filler plasmid as a function of mass of Cas13d plasmid transfected. Measurements relative to NT gRNA. Error bars indicate standard deviation. n=3 transfections per condition. Logistic functions fit to the data are shown as solid lines. **p<0.01, two-tailed Student’s *t* test. n.s.: not significant, p>0.05. **H** Splicing inclusion of the *MBNL1* exon 5 minigene measured by RT-PCR and capillary electrophoresis. The mass of Cas13d plasmid transfected matches the condition highlighted in (G) (box with dashed lines). Error bars indicate standard deviation. n=3, all conditions. *p<0.05, **p<0.01, ***p<0.001, two-tailed Student’s *t* test.

We constructed a differential equation model to describe the dynamics of GENO, and we proved that GENO strictly reduces Cas13d mRNA expression and binary complex production compared to an unregulated reference design for all possible transcription, translation, and crRNA processing rates (see Supplemental Note). To estimate how efficiently Cas13d:gRNA binary complex concentration is reduced by GENO, we performed simulations of the dynamical model across broad ranges of plausible biochemical parameters (Supplemental Fig. 5). From this analysis, we predicted that GENO robustly reduces binary complex concentration at equilibrium across a wide range of transcription rates of the Cas13d promoter (Fig. 4C), yet this reduction begins to break down at very high expression levels, at which the concentration in the unregulated design plateaus due to limited gRNA availability. We also predicted that the autoregulation efficiency (*η*_GENO_, defined as the difference of unity and the ratio of equilibrium binary complex concentration in GENO vs. unregulated designs, see Supplemental Note) increases with Cas13d translation rate (Fig. 4D). Interestingly, we found a more complex relationship between autoregulation efficiency and crRNA processing rate, exhibiting a positive correlation at high translation rates and a negative correlation at low translation rates (Fig. 4D). Overall, these insights from analytical solutions and dynamical simulations suggest that GENO is a simple and robust approach to regulate Cas13d expression in mammalian cells.

To experimentally determine if GENO reduces Cas13d expression, we transfected plasmids encoding GENO-regulated and unregulated Cas13d into HeLa cells and measured Cas13d protein expression by Western blot (Fig. 4E). In this context, we found that GENO reduced Cas13d protein concentration by 76% (Fig. 4F; p<0.001, two-tailed Student’s *t* test), validating its utility as a strategy for regulating Cas13d expression.

### Negative autoregulation reduces collateral activity while maintaining partial on-target rescue

To confirm that regulation of Cas13d expression with GENO reduces the extent of collateral activity upon targeting CUG repeat RNA, we transfected plasmids encoding GENO-regulated or unregulated Cas13d and gRNA into HeLa cells along with CUG_480_ target and mCherry plasmids, and we measured relative mCherry fluorescence between CUG-targeting and non-targeting conditions. We transfected the Cas13d plasmids across a wide range of concentrations (0.5 to 50 ng per transfection) to estimate the window of therapeutic benefit provided by GENO. With unregulated Cas13d expression, we observed strong reductions in mCherry fluorescence at all plasmid concentrations tested (Fig. 4G; p<0.01, two-tailed Student’s *t* test), underscoring the need for a solution to collateral activity for repeat-targeting Cas13d therapies. We found that GENO substantially reduced collateral activity, to the point where it was statistically undetectable in our assay for all but the two highest plasmid concentrations (p>0.05, two-tailed Student’s *t* test). By fitting logistic functions to these data, we determined that GENO increases the IC_50_ of collateral activity by 18 times (1.6 ng to 29 ng), producing a wide window in which collateral activity is improved.

To evaluate if on-target activity at CUG_n_ RNA is maintained upon GENO regulation, we chose a Cas13d plasmid concentration (5 ng per transfection) at which collateral activity was not detected with GENO, and we measured MBNL alternative splicing activity using the *MBNL1* exon 5 minigene assay. We found that GENO-regulated Cas13d partially rescued MBNL-mediated splicing of this event (Fig. 4H; 36% of rescue observed without regulation, p<0.05, two-tailed Student’s *t* test). Overall, we found that GENO preserved on-target activity of Cas13d at CUG repeat RNA while minimizing collateral activity, validating the general concept of autoregulation to control Cas13d expression. Further experimentation and optimization will be needed to maximize desired on-target knockdown while maintaining collateral effects below a tolerable threshold.

### Negative autoregulation reduces AAV-delivered Cas13d expression in human DM1 myoblasts

As a proof-of-concept for applying negative autoregulation to an AAV-Cas13d therapy, we sought to investigate if GENO efficiently regulates expression when delivering Cas13d by AAV and if GENO-regulated Cas13d reduces CUG_n_ RNA accumulation in patient-derived DM1 myoblasts. We synthesized recombinant AAV6 vectors carrying viral genomes encoding unregulated or GENO-regulated Cas13d driven by a CMV promoter with either CUG-1 or NT gRNAs. Despite multiple attempts, we were unable to package the unregulated CUG-targeting Cas13d design, possibly as a result of toxicity of overexpressed CUG-targeting Cas13d during virus production. Thus, subsequent experiments focused on GENO-regulated Cas13d paired with CUG-1 or NT gRNAs, and the unregulated Cas13d paired with the NT gRNA.

We first treated undifferentiated DM1 myoblasts with AAV for 6 days and performed a Western blot to measure Cas13d protein expression (Fig. 5A). We found that GENO reduced Cas13d protein production by 87% (p<0.05, two-tailed Student’s *t* test, n=3). To probe RNA and protein expression at the single-cell level, we simultaneously performed hybridization chain reaction FISH (HCR FISH) (Choi et al., 2018) to image single molecules of Cas13d mRNA and IF to measure Cas13d protein (Fig. 5B), and we quantitated confocal microscopy images using automated image analysis techniques (see Methods). In images taken at 40x magnification, we detected a mean of 156 diffraction-limited HCR FISH spots per nucleus in cells treated with AAV containing unregulated Cas13d and NT gRNA and 1.3 spots per nucleus in PBS-treated cells (Supplemental Fig. 6A, 6B), highlighting the sensitivity and specificity of this approach for measuring Cas13d RNA expression. Additionally, all nuclei in AAV treatment conditions (n=142) contained >5 HCR FISH spots, indicating nearly 100% efficiency of AAV transduction.

**Figure 5.**
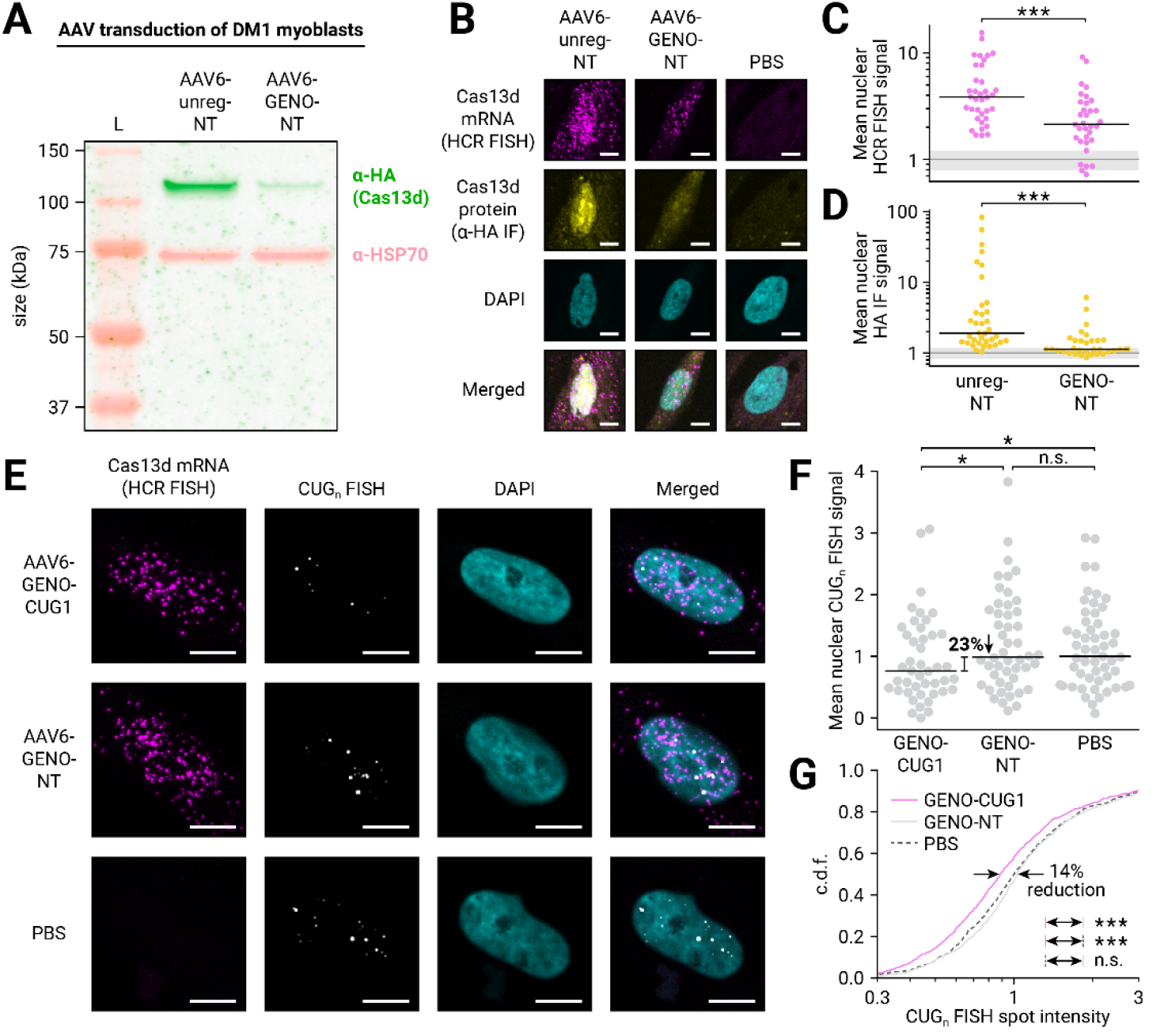
AAV-delivered autoregulated Cas13d reduces CUG_n_ RNA accumulation in human DM1 myoblasts. **A** Fluorescent Western blot comparing Cas13d protein expression from AAV transduction of patient-derived DM1 myoblasts. Cas13d is visualized with α-HA antibody (green), and HSP70 is stained as a loading control (pink). Protein ladder is shown (L). **B** Representative images from confocal microscopy of AAV-treated DM1 myoblasts stained for Cas13d mRNA (HCR FISH, magenta) and protein (α-HA IF, yellow). DAPI was used to visualize the nucleus (cyan). Images taken at 20x magnification. Scale bars 10 μm. **C** Mean nuclear intensity of Cas13d HCR FISH across nuclei in unregulated and GENO-regulated non-targeting conditions. Dots represent individual nuclei, black line indicates median. n>33 nuclei per condition, >3 images per condition. ***p<0.001, two-sided Mann-Whitney *U* test. Grey line indicates mean baseline nuclear FISH signal in PBS-treated myoblasts and grey shaded region indicates standard deviation, n=26 nuclei, 3 images. **D** Mean nuclear intensity of α-HA IF across nuclei in unregulated and GENO-regulated non-targeting conditions. Dots represent individual nuclei, black line indicates median. n>33 nuclei per condition, >3 images per condition. ***p<0.001, two-sided Mann-Whitney *U* test. Grey line indicates mean baseline nuclear FISH signal in PBS-treated myoblasts and grey shaded region indicates standard deviation, n=26 nuclei, 3 images. **E** Representative images from confocal microscopy of AAV-treated DM1 myoblasts stained for Cas13d mRNA (HCR FISH, magenta) and CUG_n_ RNA (CAG_10_ FISH probe, greyscale). DAPI was used to visualize the nucleus (cyan). Images taken at 40x magnification. Scale bars 10 μm. **F** Mean nuclear intensity of CUG_n_ FISH across nuclei in GENO-regulated targeting and non-targeting conditions. Dots represent individual nuclei, black line indicates median. n>43 nuclei per condition, 21 images per condition. *p<0.05, one-sided Mann-Whitney *U* test. n.s.: not significant, p>0.05. **G** Cumulative distribution function (c.d.f.) plot of the diffraction-limited FISH spot intensity of individual CUG_n_ RNA foci in GENO-regulated targeting (magenta, solid) and non-targeting (grey, solid) conditions. PBS-treated condition is also shown (grey, dashed). ***p<0.001, two-sided Mann-Whitney *U* test. n.s.: not significant, p>0.05.

We observed that GENO led to reduced Cas13d mRNA and protein expression compared to unregulated Cas13d when both were paired with NT gRNAs (Fig. 5B). By quantifying mean HCR FISH intensity in each nucleus imaged at 20x magnification, we found that GENO reduced median Cas13d mRNA expression by 60% after baseline subtraction of the median in the PBS-treated control (Fig. 5C, n>33 nuclei per condition, p<0.001, two-sided Mann-Whitney *U* test). Similarly, by performing the same analysis on α-HA IF, we found an 83% reduction in median Cas13d protein expression with GENO (Fig. 5D, n>33 nuclei per condition, p<0.001, two-sided Mann-Whitney *U* test). These results illustrate that negative autoregulation can reduce Cas13d expression in human myoblasts and that GENO-regulated Cas13d can be easily delivered by AAV.

### Autoregulated Cas13d reduces CUG_n_ RNA accumulation in human DM1 myoblasts

To evaluate if GENO-regulated Cas13d can reduce nuclear CUG_n_ RNA foci in patient-derived cells, we transduced DM1 myoblasts with AAV for 6 days, and we simultaneously performed FISH with a fluorescent CAG_10_ probe to detect CUG_n_ RNAs from the expanded *DMPK* allele and HCR FISH for Cas13d mRNA to mark transduced nuclei (Fig. 5E). We collected confocal images at 40x magnification to visualize discrete diffraction-limited spots in both channels, and we detected FISH spots and quantified fluorescence intensities using automated image analysis (see Methods).

Upon treatment with GENO-regulated Cas13d and CUG-1 gRNA, we found a modest 23% reduction in median CUG_n_ RNA signal per nucleus (Fig. 5F, n>43 nuclei, p<0.05, one-sided Mann-Whitney *U* test) relative to NT gRNA and PBS-treated controls, with no significant difference between these controls (p>0.05, n>43 nuclei, two-sided Mann-Whitney U test). Although absolute foci number was not significantly different (Supplemental Fig. 6D, p>0.05, two-sided Mann-Whitney *U* test), we did observe a 14% reduction in median foci intensity (Fig. 5G, n>745 spots per condition, p<0.001, two-sided Mann-Whitney *U* test). Interestingly, we did not observe a similar reduction in Cas13d HCR FISH intensity between targeting and non-targeting conditions (Supplemental Fig. 6C), suggesting that the extent of collateral activity of GENO-regulated Cas13d in DM1 myoblasts may be low enough to preserve RNA homeostasis. These results suggest that GENO-regulated Cas13d modestly reduces accumulation of expanded CUG_n_ RNAs in DM1 patient-derived cells by cleaving and dispersing multimeric CUG_n_ RNA structures and initiating RNA decay.

## DISCUSSION

The extent of Cas13 collateral activity in mammalian cells is controversial; many groups have found it to be negligible in eukaryotic cells (Abudayyeh et al., 2017; Huynh et al., 2020; Konermann et al., 2018; Kushawah et al., 2020), while evidence has begun to emerge of *trans* cleavage in certain contexts (Özcan et al., 2021; Wang et al., 2021a, 2019b; Xu et al., 2021). We observed stark collateral activity of Cas13d in mammalian cells using fluorescence, cell viability, and whole transcriptome assays, yet there are several plausible reasons why we have observed intense effects while others have not. For one, non-specific RNase activity may go undetected by assays that identify relative changes to transcript abundances, such as RNA-seq, unless it is strong enough to cause upregulation of stress response and apoptosis pathways. Additionally, we observed a positive relationship between the total protospacer load and the extent of collateral activity, with collateral activity being the strongest when targeting overexpressed repetitive RNAs (Fig. 2C-G), weaker when targeting overexpressed transgenic RNAs at unique protospacer sequences (Fig. 2H), still weaker when targeting highly expressed endogenous genes (Fig. 3C), and undetectable when targeting lowly expressed genes (Fig. 3C). Therefore, we would not necessarily expect reports of collateral activity with these or similar assays if moderately or lowly expressed genes are targeted using a single gRNA against a unique protospacer—a particularly common use case for Cas13d. Importantly, our observations do not preclude any use of Cas13d for RNA knockdown; rather, we suggest that researchers thoroughly assess bystander RNA cleavage when targeting any gene with Cas13d in mammalian cells. Furthermore, we propose a solution for targeting repeat expansion RNAs, a setting that presents both extreme challenges and unique opportunities for mitigating toxic effects of collateral activity.

Notably, some of Cas13d’s most successful applications in mammalian cells have been cases in which collateral activity may synergistically align with experimental goals rather than detract from or confound them. For example, Cas13d targeting cancer-specific transcripts can induce apoptosis of bladder and pancreatic cancer cells *in vitro* and halt tumor progression in xenograft models (Jiang et al., 2020; Li et al., 2021a; Zhuang et al., 2021). In this context, extensive collateral activity would likely exacerbate cancer-specific toxicity by interfering with critical cellular processes. Cas13d has also been used to convert glia to neurons *in vivo* by targeting the RNA-binding protein gene *Ptbp1* (Zhou et al., 2020) (although these findings are actively disputed (Wang et al., 2021b)). An intriguing hypothesis is that collateral activity may be useful in promoting differentiation and cell type conversion by clearing mRNAs globally, inducing a transcriptomic reset that allows the cell to quickly reestablish a new stable gene expression state in response to on-target knockdown. Thorough experimentation and direct comparison of Cas13d with RNA-targeting approaches lacking collateral activity (such as shRNA) are required to discern whether collateral activity plays an important advantageous role in these applications.

One group has observed collateral activity of Cas13a in some cell lines but not others and has proposed its application to specifically kill cancer cells (Wang et al., 2019b). In contrast, we observed similar levels of collateral activity when targeting exogenous repeats with Cas13d in all cell types tested, including both human- and mouse-derived cell lines as well as cancerous and noncancerous lines. Still, this does not preclude the involvement of cell type and cell state on the extent of collateral activity more generally. Differential expression of Cas13, gRNA, target RNA, and other factors such as RNA degradation and proteostasis machinery may influence the rate of ternary complex dissociation, which likely determines the expected number of cleaved bystander RNAs per target RNA. We therefore propose that collateral activity of Cas13d in mammalian cells is a general phenomenon and not strictly cell-type-specific, yet the extent of collateral activity is a complex function of many variables, many of which vary widely among cell types. Nevertheless, therapeutic application of collateral activity to induce apoptosis in specific cells, e.g. by targeting cancer-specific mutant transcripts (Jiang et al., 2020) or viral RNAs (Wang et al., 2021a), remains an enticing concept.

To combat the detrimental effects of collateral activity for repeat expansion disease therapies, we have developed GENO, a straightforward negative autoregulation approach that repurposes the innate crRNA processing functionality of Cas13. A fundamental regulatory motif in natural and synthetic gene networks (Alon, 2006), negative autoregulation stabilizes gene expression temporally and intracellularly by providing a buffer against gene expression stochasticity (Becskei and Serrano, 2000). Autoregulation also decreases the response time to reach equilibrium after perturbations (Rosenfeld et al., 2002), including AAV transduction and initial cargo expression. Nearly half of transcription factors in *Escherichia coli* (Shen-Orr et al., 2002) and many RBPs in humans (Pervouchine et al., 2019) natively exhibit negative autoregulation, highlighting its evolutionary importance as a method to achieve robustness in noisy biological systems. Thus, negative autoregulation may confer added benefits to gene therapies beyond reduction of cargo expression alone and should be explored further as a fundamental regulatory tool.

The simplicity of GENO also enables multiple routes for further optimization of Cas13d expression. By eliminating the RNA polymerase III promoter driving gRNA transcription, GENO reduces the sequence length of the AAV cargo, providing space for additional regulatory elements to be implemented in the final design. Additionally, our dynamical modeling predicts the performance of GENO across ranges of relevant biochemical parameters (transcription, translation, and crRNA processing rates; see Supplemental Fig. 5) and thus provides a blueprint for further tuning and optimization. In DM1 and other repeat expansion diseases (Malik et al., 2021; Sznajder and Swanson, 2019), repeat expansions often exceed thousands of repeat units in length, particularly in the most severely affected tissues (Otero et al., 2021; Thornton et al., 1994). Thus, we hypothesize that very low expression of an RNA-targeting therapy would reduce off-target interactions with transcripts containing short repeats while maintaining high probability of on-target cleavage. With this insight in mind, our simulations suggest that optimal autoregulation (with weak but consistent Cas13d expression) is generally achieved when transcription rate is low but translation and crRNA processing rates are high. Nevertheless, the optimal binary complex concentration will likely depend on the specific application of GENO to different diseases and target tissues. Transcription and translation rates can be tuned by adjusting promoter and Kozak sequences, and single-base mutations in the direct repeat region of the pre-crRNA have been shown to affect crRNA processing rate (Zhang et al., 2019), providing a potential engineering mechanism to adjust this parameter as well. We believe that this theoretical characterization lays groundwork for extending GENO to Cas13-based therapies for many other repeat expansion diseases.

Crucially, the feasibility of a negative autoregulation approach to mitigate collateral activity relies on an expectation of nonlinearity between *cis* and *trans* knockdown efficiency caused by the presence of many protospacer sequences on a repetitive target RNA (Fig. 4A). As a result, to mitigate collateral activity at targets with unique protospacers, other solutions may be needed that reduce the rate of *trans* cleavage for each activated Cas13d ternary complex while maintaining strong knockdown of the intended target. Comprehensive characterization of intracellular collateral activity across Cas13 orthologs, especially small and truncated variants that are compatible with AAV delivery (Kannan et al., 2021; Xu et al., 2021), may reveal effectors with better selectivity than RfxCas13d, with the caveat that these differences may be target- and context-dependent. The newly discovered RNA-targeting CRISPR effector Cas7-11 (Özcan et al., 2021) is also a promising candidate for therapeutic knockdown approaches. Additionally, it is plausible that protein engineering strategies, including directed evolution and rational mutagenesis, may be successful in generating Cas13d variants with reduced collateral activity. As a hypothetical example, an engineered Cas13d variant with increased RNA cleavage and target dissociation rates may show improved selectivity by capitalizing on the physical proximity of the bound *cis* target RNA to the dual-R-X_4_-H catalytic site relative to bystander RNAs. Directed evolution has been applied to modify PAM requirements (Kleinstiver et al., 2015a, 2015b) and enhance specificity of Cas9 (Cerchione et al., 2020; Vakulskas et al., 2018) and to improve Cas13a stability in eukaryotic cells (Charles et al., 2021). We believe this approach is a promising strategy to improve collateral activity of Cas13d for more general applications.

In some situations, it may be possible to achieve similar therapeutic goals using nuclease-inactive dCas13d, for example by blocking translation initiation (Charles et al., 2021) or interfering with RNA-binding proteins. We had considered applying dCas13d to displace MBNL proteins from CUG_n_ RNA and rescue splicing phenotypes; however, although we observed dCas13d enriched at RNA foci in the nucleus (Supplemental Fig. 1C), it did not displace MBNL sufficiently to detect an increase in MBNL-dependent splicing (Fig. 1G), and much of the dCas13d in the nucleoplasm remained unbound (Supplemental Fig. 1B). It is important to note that these observations may result from performing these experiments in an overexpression context, where CUG_480_ RNA may fully deplete MBNL with many sites along the RNA remaining accessible. However, these results may also indicate that the affinity of the dCas13d binary complex to CUG_n_ RNA is insufficient to adequately displace MBNL. Indeed, the myriad intra- and intermolecular structures explored by repeat expansion RNAs and RNA-binding proteins (Zhang and Ashizawa, 2017) may thermodynamically inhibit binding of dCas13. It also remains plausible that the target affinity of dCas13 is weak in general; in support of this, a recent study showed that dCas13d fused to APEX2 was unable to bind the human telomerase RNA (hTR) unless also fused to a double-stranded RNA-binding domain (dsRBD) to enhance RNA binding affinity, even for gRNAs that produced strong knockdown of hTR by active Cas13d (Han et al., 2020). Although notable successes in RNA imaging (Abudayyeh et al., 2017; Yang et al., 2019), RNA editing (Cox et al., 2017; Kannan et al., 2021; Xu et al., 2021), and alternative splicing modulation (Du et al., 2020; Konermann et al., 2018) appear to contradict this hypothesis, it is possible that globally weak binding affinity can be compensated by overexpression of dCas13 in many situations. Interestingly, in the context of CRISPR-mediated phage immunity, the altruistic and self-preserving goals proposed to explain the evolutionary development of Cas13 systems (Abudayyeh et al., 2016; Meeske et al., 2019) would seem to pressure Cas13 target affinity in opposing directions: a slow target dissociation rate, while further inhibiting replication and release of phage through prolonged *cis* and *trans* RNA cleavage, might counterproductively reduce the chance of survival after depletion of host RNAs. Further experimentation is needed to characterize the binding affinity of dCas13 orthologs across a wide range of target RNA sequences and to deeply understand the role that target affinity plays in the evolutionary processes driven by class 2 type VI CRISPR systems.

Finally, although CRISPR systems are being rapidly developed as therapeutic platforms for human disease, many other challenges lie ahead. Efficient and safe delivery of large nucleic acid therapies to specific tissues remains a significant hurdle, although recent advances in AAV capsid engineering (such as MyoAAV for delivery to skeletal muscle (Tabebordbar et al., 2021)) and nonviral delivery approaches (Mitchell et al., 2021) inspire confidence that these obstacles will be overcome in the near future. Additionally, immunogenicity of non-human proteins, including CRISPR effectors in particular (Hakim et al., 2021), may stand in the way of long-term expression of Cas proteins for therapeutic use. Nevertheless, our work here to specifically characterize and minimize collateral activity of Cas13d in repeat expansion diseases outlines general principles for how to select properties of optimal therapeutic cargoes, which will ultimately depend on application and disease context. Overall, this work underscores the need to develop universally robust regulatory circuits for gene therapies that enable precise targeting of narrow therapeutic windows, particularly in situations where uncontrolled overexpression of the therapeutic is toxic. We believe that GENO serves as an example for the design of simple and compact engineering solutions to optimize potency and safety of gene therapies for a wide range of human diseases.

## ACKNOWLEDGMENTS

We are grateful to all members of the Wang Lab for their continuous support and collaborative environment, and in particular Ona McConnell and Ryan Hildebrandt for their assistance with cell culture experiments. We would like to thank the University of Florida Interdisciplinary Center for Biotechnology Research for providing flow cytometry services. We also thank Yuxi Ai and Prof. Jeremy Wilusz (University of Pennsylvania) for our collaborative discussions on Cas13 collateral activity. Finally, we would like to thank Geno the Yorkshire terrier, the eponym for our autoregulation approach. This work was supported by NIH R01 AG058636 awarded to E.T.W. C.P.K. is supported by the NSF Graduate Research Fellowship.

---

**Supplemental Figure 1.**
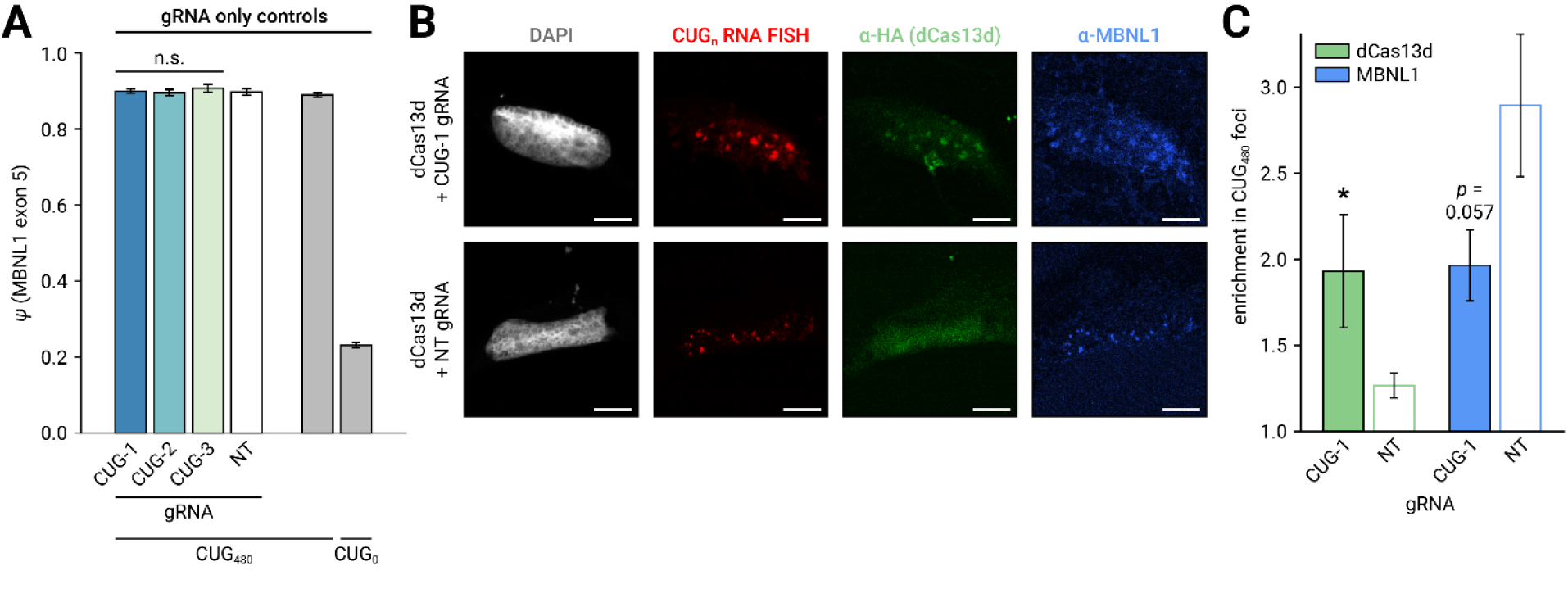
gRNA only controls for splicing assay and dCas13d/MBNL1 competition. **A** *MBNL1* exon 5 minigene splicing assay after transfection of HeLa cells with gRNA and target plasmids, in the absence of Cas13d. n=3 transfections per condition. n.s.: not significant, p>0.05, two-tailed Student’s *t* test. **B** Simultaneous FISH/IF of transfected HeLa cells for CUG_480_ RNA (CAG_10_ FISH probe, red), dCas13d (α-HA IF, green), and MBNL1 (α-MBNL1 IF, blue). Nuclei stained with DAPI (white). Scale bars 10 μm. **C** Quantification of colocalization of dCas13d and MBNL1 IF signal with nuclear CUG_480_ RNA foci in FISH/IF experiment. n=4 nuclei per condition. *p<0.05, two-sided Mann-Whitney *U* test.

**Supplemental Figure 2.**
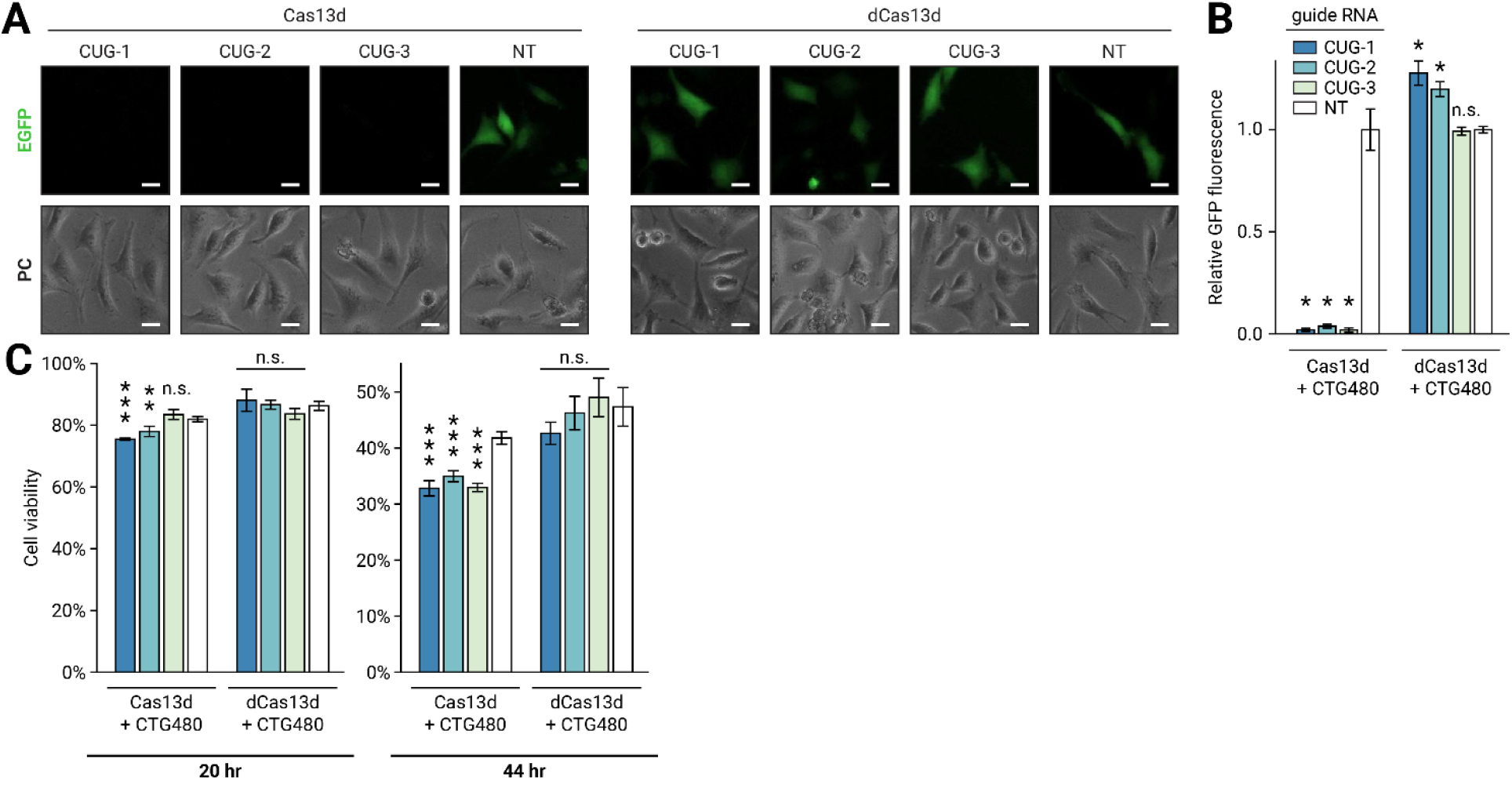
CUG-targeted Cas13d suppresses EGFP expression and reduces cell viability. **A** Visualization of unfused EGFP marker on Cas13d plasmid 20 hr after transfection with Cas13d, gRNA, and CUG_480_ target plasmids. PC: phase contrast. Scale bars 20 μm. **B** Quantification of EGFP expression by plate reader 20 hr after transfection. n=3 transfections per condition. Error bars indicate standard deviation. *p<0.05, two-tailed Student’s *t* test. n.s.: not significant, p>0.05. **C** Resazurin cell viability assay performed 20 hr and 44 hr after transfection with Cas13d, gRNA, and CUG_480_ target plasmids. n=5 transfections per condition. Error bars indicate standard deviation. *p<0.05, **p<0.01, ***p<0.001, two-tailed Student’s *t* test. n.s.: not significant, p>0.05.

**Supplemental Figure 3.**
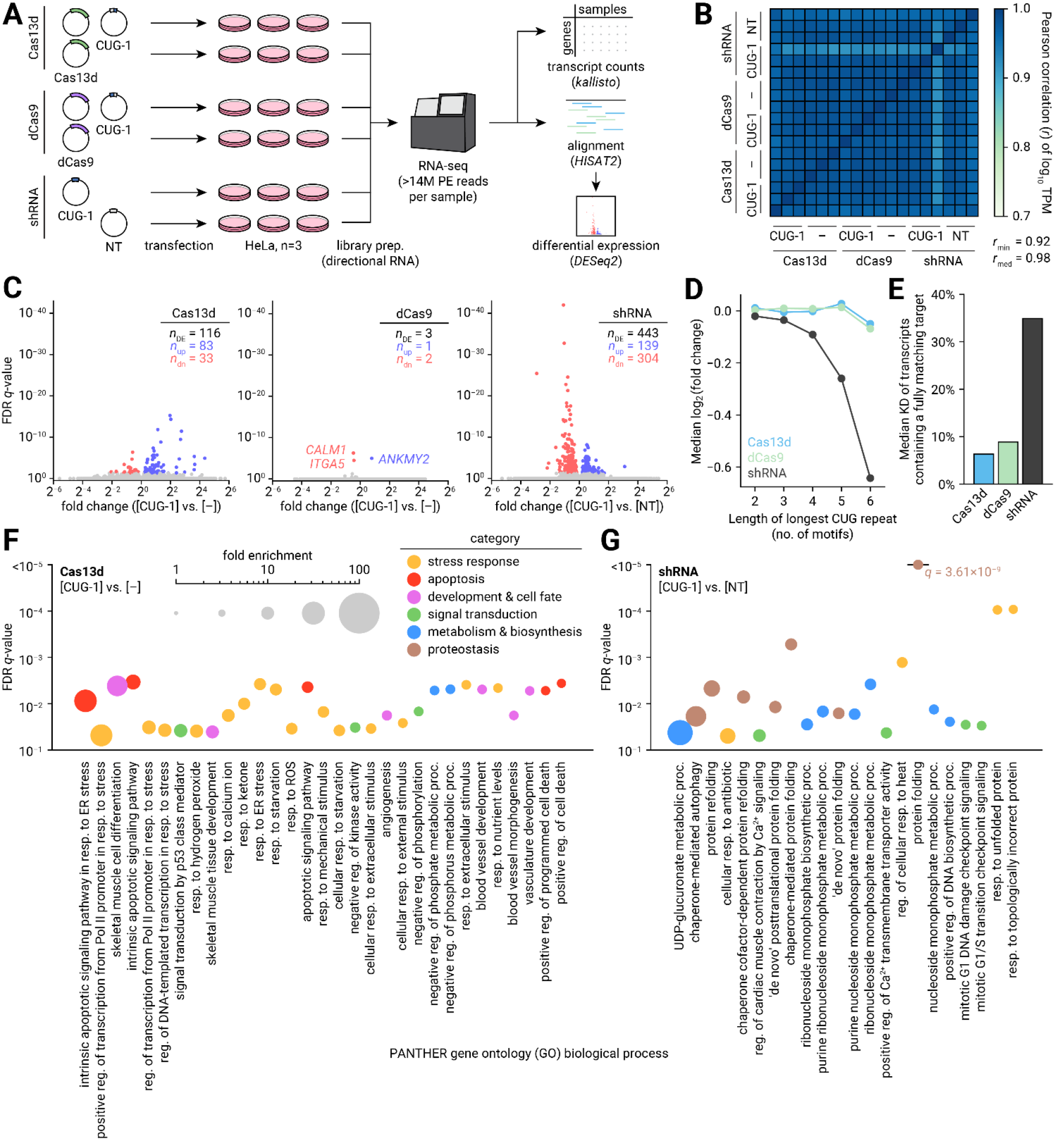
CUG-targeted Cas13d upregulates stress response and apoptosis pathways in HeLa. **A** Description of RNA-seq experiment to assess transcriptomic changes induced by Cas13d. HeLa cells were transfected with Cas13d, dCas9, or shRNA in CUG-targeting or non-targeting conditions and incubated for 3 days prior to RNA extraction, library preparation, and sequencing. n=3 transfections per condition. Data were processed using kallisto, HISAT2, and DESeq2 for alignment and differential expression (DE) analysis. **B** Heatmap of correlation coefficients of log_10_ TPM between sequencing libraries. **C** Volcano plots of DESeq2 false discovery rate (FDR)-corrected q-value vs. fold change in targeting and non-targeting conditions. DE genes (FDR q<0.05) are highlighted in red (downregulated) or blue (upregulated). **D** Plot of median fold change of transcripts in targeting and non-targeting conditions, binned by maximum CUG repeat length within the transcript in the human reference genome. **E** Median knockdown between targeting and non-targeting conditions of all transcripts containing a CUG repeat as long as or longer than the length of the Cas13d spacer (22 nt). **F** PANTHER gene ontology (GO) analysis of biological processes enriched in the DE genes between CUG-targeting and non-targeting Cas13d conditions. Enriched processes are defined as processes with a ratio of observed to expected genes >5 and FDR q<0.05. For each process, FDR q is plotted on the vertical axis and enrichment is indicated by circle area. Color indicates classification into functional categories. **G** PANTHER GO analysis of processes enriched in DE genes between CUG-targeting and non-targeting shRNA conditions.

**Supplemental Figure 4.**
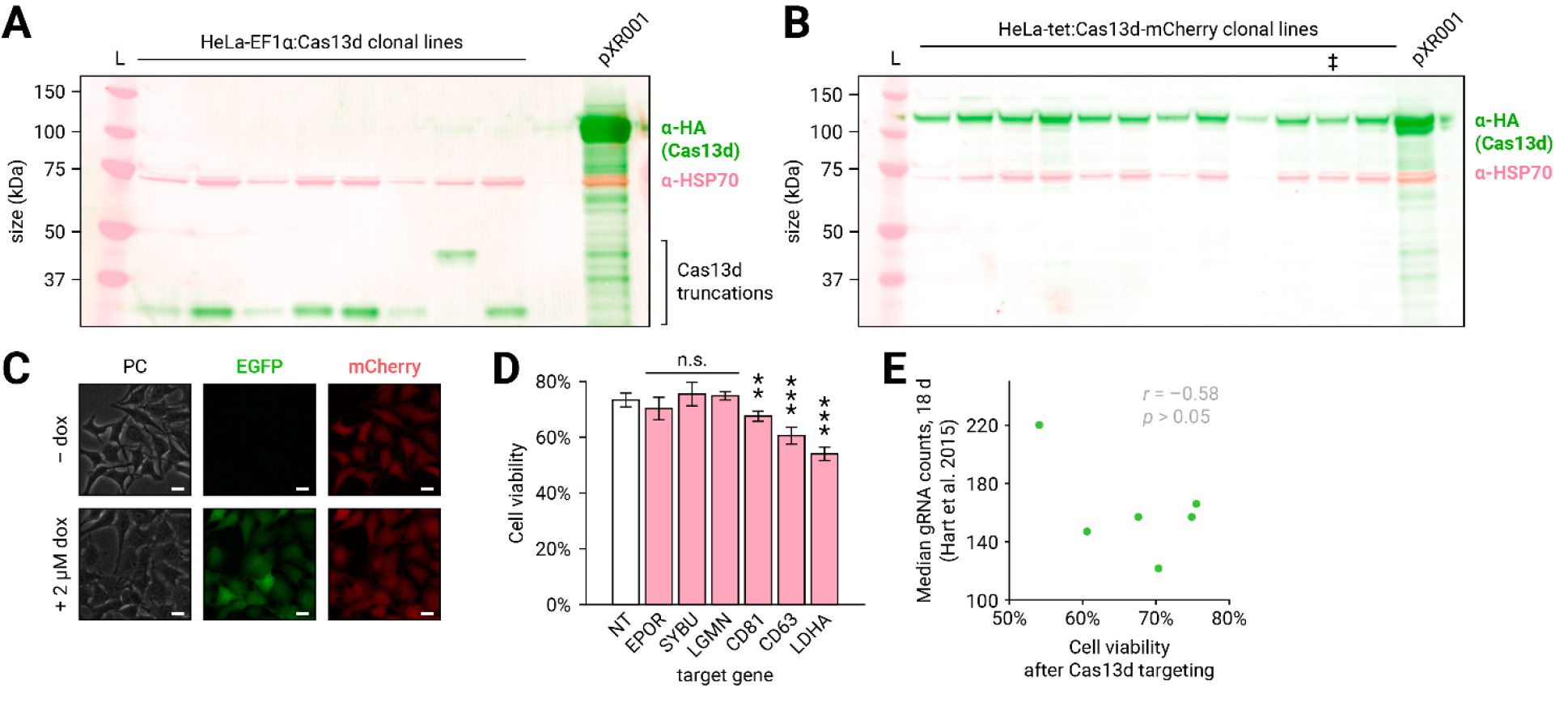
Development of HeLa-tet:Cas13d-mCherry cell line and cell viability upon targeting endogenous genes. **A** Fluorescent Western blot of protein extracted from clonal HeLa cell lines after treatment with lentivirus encoding Cas13d-T2A-EGFP under the constitutive EF1α promoter. Blot stained with α-HA (green) and α-HSP70 (red) primary antibodies. Expected MW of Cas13d is 117 kDa, lower bands indicate truncations of Cas13d that retained expression of the downstream EGFP marker. L: protein ladder, pXR001: transient transfection of Cas13d plasmid in HeLa. **B** Fluorescent Western blot of protein extracted from clonal HeLa cell lines after integration of constitutive mCherry and tetracycline-inducible Cas13d-T2A-EGFP. Expression induced with 2 uM doxycycline for 44 hr prior to protein extraction. Blot stained with α-HA (green) and α-HSP70 (red) primary antibodies. ‡ indicates the clone chosen for subsequent experiments. L: protein ladder, pXR001: transient transfection of Cas13d plasmid in HeLa. **C** Visualization of EGFP and mCherry before and after 44 hr doxycycline treatment by fluorescence microscopy. PC: phase contrast. Scale bars 20 μm. **D** Resazurin cell viability assay of HeLa-tet:Cas13d-mCherry cells transfected with plasmids encoding gRNAs targeting endogenous genes and induced with 2 uM doxycycline for 44 hr. n=5 transfections per condition. Error bars indicate standard deviation. *p<0.05, **p<0.01, ***p<0.001, two-tailed Student’s *t* test. n.s.: not significant, p>0.05. **E** Comparison of cell viability measured 44 hr after Cas13d expression with gRNA targeting endogenous genes vs. depletion of gRNAs targeting the same genes in a CRISPR essentiality screen in HeLa (Hart et al., 2015). Pearson correlation coefficient is shown. p>0.05, beta distribution c.d.f.

**Supplemental Figure 5.**
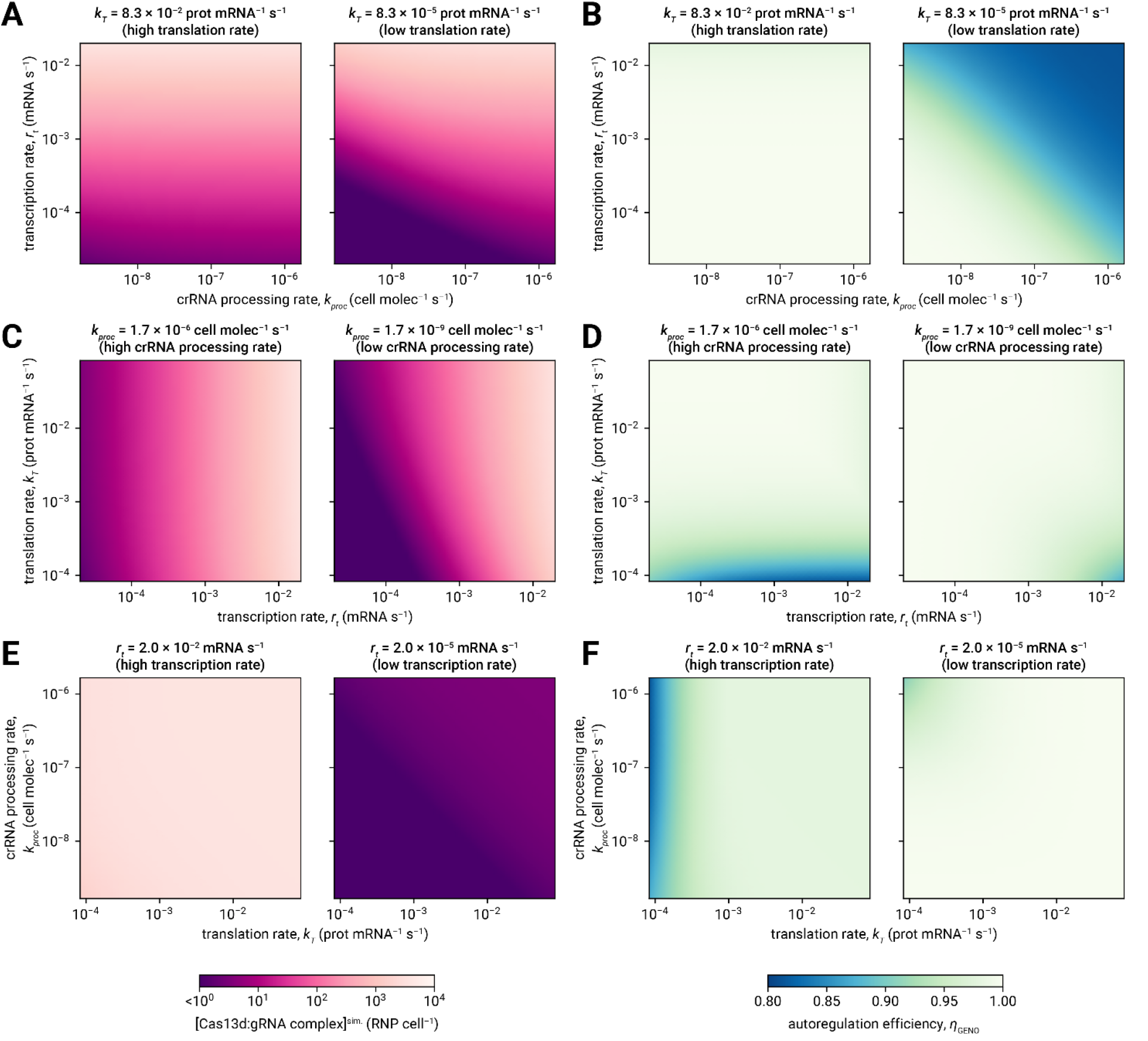
Prediction of Cas13d binary complex concentration and autoregulation efficiency from simulation of GENO dynamics. **A** Equilibrium Cas13d binary complex concentration as a function of transcription rate and crRNA processing rate, for high (left) and low (right) translation rate. **B** Equilibrium binary complex concentration as a function of translation rate and transcription rate, for high (left) and low (right) crRNA processing rate. **C** Equilibrium binary complex concentration as a function of crRNA processing rate and translation rate, for high (left) and low (right) transcription rate. **D** Autoregulation efficiency (*η*_GENO_, defined in Supplemental Note) as a function of transcription rate and crRNA processing rate, for high (left) and low (right) translation rate. **E** *η*_GENO_ as a function of translation rate and transcription rate, for high (left) and low (right) crRNA processing rate. **F** *η*_GENO_ as a function of crRNA processing rate and translation rate, for high (left) and low (right) transcription rate.

**Supplemental Figure 6.**
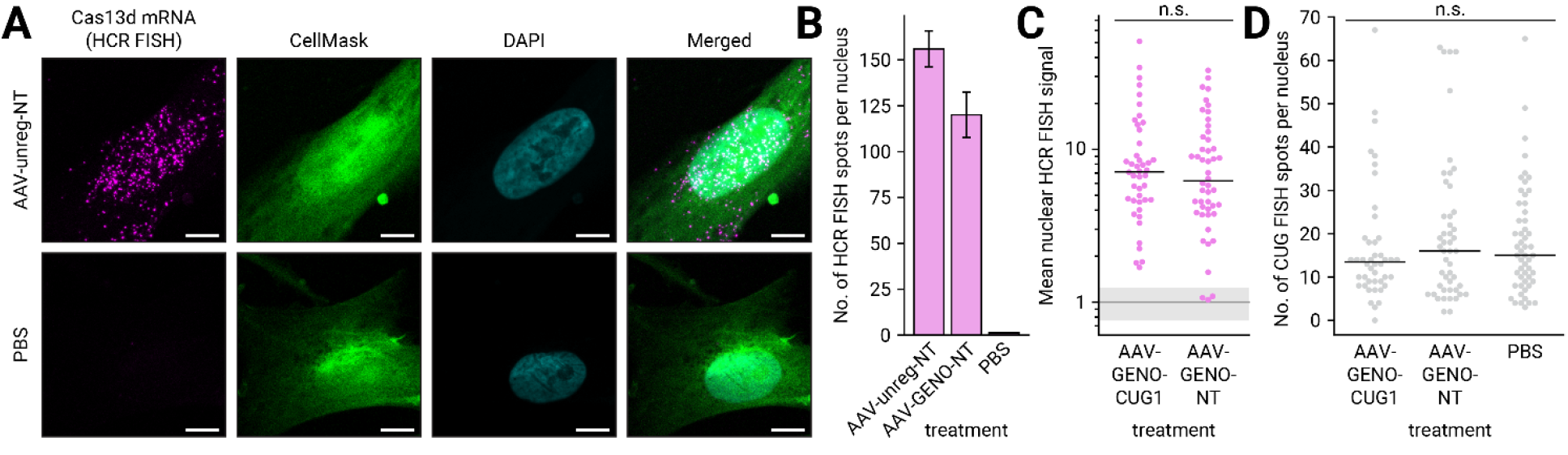
Detection of diffraction-limited spots in Cas13d HCR FISH and CUG_n_ FISH images of AAV-treated DM1 myoblasts. **A** Representative images at 40x magnification of DM1 myoblasts stained for Cas13d mRNA (HCR FISH, magenta) after treatment with AAV for 6 days. Cytoplasm (CellMask, green) and nuclei (DAPI, cyan) are also labeled. Scale bars 10 μm. **B** Mean number of Cas13d HCR FISH spots detected in nuclei after AAV treatment. Error bars indicate SEM. **C** Mean nuclear intensity of Cas13d HCR FISH across nuclei in GENO-regulated targeting and non-targeting conditions. Dots represent individual nuclei, black line indicates median. n>43 nuclei per condition, 21 images per condition. n.s.: not significant, p>0.05, two-sided Mann-Whitney *U* test. Grey line indicates mean baseline nuclear FISH signal in PBS-treated myoblasts and grey shaded region indicates standard deviation, n=53 nuclei, 21 images. **D** Number of CUG_n_ FISH spots (RNA foci) detected in each nucleus after AAV treatment. Dots represent individual nuclei, black line indicates median. n>43 nuclei per condition, 21 images per condition. n.s.: not significant, p>0.05, two-sided Mann-Whitney *U* test.

## METHODS

### Plasmids and molecular cloning

Plasmids encoding NLS-RfxCas13d-NLS-HA-T2A-EGFP (pXR001) and NLS-dRfxCas13d-NLS-HA-T2A-EGFP (pXR002) under the EF1α promoter were purchased from Addgene (#109049 and #109050, respectively), as was a plasmid encoding a gRNA cloning cassette driven by the U6 promoter (pXR003, #109053) (Konermann et al., 2018). All plasmids in this study were propagated in NEB Stable chemically competent *E. coli* (New England Biolabs (NEB), #C3040) at 30°C and purified using the Zyppy Plasmid Miniprep kit (Zymo Research, #D4036) or ZymoPURE II Plasmid Midiprep kit (Zymo Research, #D4200). Spacer sequences were cloned into pXR003 by BbsI digestion and Gibson assembly (NEB, #E2611) with synthesized DNA duplexes containing the spacer sequence flanked by 19 bp homology arms (Integrated DNA Technologies (IDT)). Plasmids encoding nuclease-inactive *Streptococcus pyogenes* Cas9 (dCas9) and gRNAs were utilized from a previous study (Pinto et al., 2017). To match the expression context of Cas13d, dCas9 was cloned into the pXR001 vector by removing Cas13d with BsiWI and NheI and inserting a PCR-amplified dCas9 amplicon by Gibson assembly. CUG-targeting and non-targeting short hairpin RNAs (shRNAs) matching the corresponding Cas13d spacer sequences were cloned into the pLKO.1 vector (Addgene, #10878) by AgeI and EcoRI digestion and ligation of 5’-phosphorylated DNA duplexes using T4 DNA ligase (NEB, #M0202). Target plasmids expressing 0 and 480 CTG repeats in the context of *DMPK* exons 11-15 (DMPKS and DT480, respectively) were gifted by Tom Cooper (Baylor College of Medicine).

To investigate collateral activity, a plasmid expressing mCherry under the CMV promoter (pmCherry) was cloned by removing EGFP from pEGFP-C1 (Clontech) with AgeI and BglII and assembling the vector with a synthesized mCherry gene fragment (IDT) using In-Fusion cloning (Takara Bio, #102518). pB-Tet-Cas13d was cloned using Gibson assembly by inserting a fragment PCR-amplified from pXR001 containing NLS-RfxCas13d-NLS-HA-T2A-EGFP into a piggyBac transposon vector (Cadiñanos and Bradley, 2007) expressing the insert in a Tet-On cassette and constitutively expressing puromycin acetyltransferase (pac) and reverse tetracycline-controlled transactivator (rtTA) (Gossen et al., 1995). pB-mCherry was cloned using Gibson assembly by inserting a PCR-amplified mCherry gene from a synthetic gene fragment (IDT) into a piggyBac transposon vector expressing the insert constitutively under an EF1α promoter and constitutively expressing a pac-thymidine kinase (TK) fusion protein.

To implement the gRNA excision negative feedback design, a synthetic DNA fragment containing CUG-1 or NT pre-crRNA (22 nt spacer flanked by two 36 nt direct repeats (DR)) was assembled by annealing two complementary oligonucleotides (IDT Ultramer) and inserted into the transcribed region of pXR001 at the KpnI site by Gibson assembly. Plasmids for recombinant AAV preparation were generated by cloning NLS-RfxCas13d-NLS-HA and gRNA (either separately driven by U6 for the unregulated design or within the 5’ UTR of the Cas13d gene as pre-crRNA for negative autoregulation) into a vector containing a CMV expression cassette flanked by AAV2 inverted terminal repeats (ITRs).

### Cell culture and transfection

HEK293, Neuro2a, HeLa, and HeLa-derived cell lines were maintained in 10% FBS growth medium (Dulbecco’s modified eagle medium (DMEM) + 10% fetal bovine serum (FBS) + 1% penicillin/streptomycin) in a humidity-controlled 5% CO_2_ incubator at 37°C. Myoblasts derived from an adult DM1 patient biopsy (DM-05 (Xia et al., 2013)) were cultured in SkGM-2 skeletal muscle cell growth medium (Lonza, #CC-3245).

For transient transfection experiments, unless otherwise stated, cells were passaged to 12-well tissue culture plates at a density of 1.5×10^5^ cells/well (3.9×10^4^ cells/cm^2^) and transfected with 500 ng plasmid DNA using 2 uL of *TransIT-X2* transfection reagent (Mirus Bio, #MIR6005) and 100 uL Opti-MEM I reduced serum medium (ThermoFisher Scientific, #31985088) according to the manufacturer’s protocol. When transfecting multiple plasmids, plasmids were mixed at equimolar concentrations unless otherwise noted. Cells were incubated with transfection reagent for 20 hr, after which the culture medium was aspirated and replaced with 10% FBS growth medium. Cells were then incubated for an additional 24 hr for 2-day experiments or 48 hr for 3-day experiments, followed by fixation or RNA isolation for further analysis.

### Fluorescence *in situ* hybridization (FISH)

Nuclear foci formed by CUG_n_ RNA were visualized by FISH using a previously described protocol (Pinto et al., 2017). Briefly, cells in 4-well coated glass chamber slides (ThermoFisher Scientific, #154917) were washed with phosphate-buffered saline (PBS) and fixed in 4% paraformaldehyde (PFA) in PBS at room temperature (RT) for 10 min, washed 3X with PBS, and permeabilized with ice-cold 70% ethanol in water and stored overnight at −20°C. Cells were washed for 30 min at 30°C in FISH wash buffer (25% formamide, 2X saline sodium citrate (SSC) in water). CAG_10_ FISH probe labeled with Alexa Fluor 594 (Biosearch Technologies, #SS151541-01) was diluted to a working concentration of 380 ng/mL in FISH hybridization buffer (100 mg/mL dextran sulfate, 1 mg/mL yeast tRNA, 2 mM ribonucleoside vanadyl complex (VRC), 200 ug/mL bovine serum albumin (BSA), 25% formamide, 2X SSC in water) and incubated with cells overnight at 30°C in a humidified chamber. The following day, the probe solution was aspirated and replaced with FISH wash buffer for 30 min at 30°C. For nuclear and whole-cell staining, 1 ng/uL DAPI and 1X CellMask Green Plasma Membrane Stain (ThermoFisher Scientific, #C37608) in PBS was added to the cells for 5 min at RT, followed by three washes with PBS of 5 min each. Slides were mounted with glass #1.5 coverslips in Fluoroshield antifade mounting medium (Sigma-Aldrich, #F6182), sealed with clear nail polish, and stored at −20°C until imaging. Widefield epifluorescence and confocal Airyscan imaging was performed on a Zeiss LSM 880 microscope with a Plan-Apochromat 40x/1.3 Oil DIC M27 objective lens. 10+ images were collected for each condition. For confocal images, Airyscan processing was performed in Zeiss ZEN software (version 2.1 SP3 FP3 black 14.0.20.201).

Image processing and FISH quantitation were performed in Python 3. 3D epifluorescence images in CZI format were separated into FISH, DAPI, and CellMask channels, and each channel was collapsed to 2D by maximum intensity projection along the z-dimension. Nuclei were segmented from the DAPI channel using Cellpose 0.0.2.0 (Stringer et al., 2021). Transfected cells were identified as those with a mean FISH intensity in the nucleus >50% higher than the background intensity, calculated as the median FISH intensity in the region outside the nuclear mask. 8+ transfection-positive nuclei were detected for each condition. The mean FISH signal was calculated for each transfected nucleus in all images for each condition. The median for each condition was calculated, and the 95% confidence interval was estimated by bootstrapping.

### Immunofluorescence (IF)

For dCas13d/MBNL1 colocalization experiments, IF was performed prior to CUG_n_ RNA FISH. Cells in 4-well chamber slides were washed with PBS and fixed in 4% PFA at RT for 10 min, washed 3X with PBS, and permeabilized with ice-cold 70% ethanol in water and stored overnight at −20°C. Cells were washed 3X with IF wash buffer (PBS + 0.1% Tween 20 (PBS-T) + 0.5 mM VRC) and incubated with IF blocking buffer (IF wash buffer + 1% BSA) for 30 min at RT. Primary antibodies (mouse α-MBNL1 (MB1a(4A8), 1:4), rabbit α-HA (C29F4, 1:1000)) were mixed in IF blocking buffer and applied to cells overnight at 4°C. The next day, cells were washed 3X with IF wash buffer. Secondary antibodies (α-mouse Alexa Fluor 647, α-rabbit Alexa Fluor 555) were added at a 1:1000 dilution to IF blocking buffer and incubated with cells for 2 hr at RT. Cells were washed 3X with PBS prior to performing FISH as described above.

### Minigene splicing assay

To measure MBNL-mediated alternative splicing activity, we used an MBNL-regulated splicing reporter minigene spanning *MBNL1* exons 4-6 cloned into the RG6 vector (RG6-MBNL1e5) (Orengo et al., 2011). RG6-MBNL1e5 was cotransfected into HeLa cells with plasmids encoding Cas13d, CUG-targeting or non-targeting gRNA, and CUG_480_ target. n=3 transfections per condition. 44 hr after transfection, RNA was extracted from cells using 300 uL TRIzol (Zymo Research, #R2050) and purified using the Direct-Zol RNA Miniprep kit (Zymo Research, #R2051) according to the manufacturer’s protocol. 100 ng RNA was reverse-transcribed into cDNA using the iScript Reverse Transcription Supermix (Bio-Rad, #1708841). Minigene isoforms were amplified by *Taq* PCR from 2 uL cDNA using forward and reverse primers complementary to the vector sequence (RG6_F, RG6_R; 54°C annealing; 28 cycles). Primer sequences are presented in Supplemental Table 4. Exon 5 inclusion ratios were quantitated from 4 uL of PCR reaction by capillary electrophoresis (Fragment Analyzer, Advanced Analytical 1.1.0.11).

### EGFP fluorescence assays

To visualize loss of EGFP expression upon Cas13d targeting, HeLa cells were transfected with plasmids encoding Cas13d and EGFP, CUG-targeting or non-targeting gRNA, and CUG480 target. n=3 transfections per condition. 20 hr after transfection, cells were imaged on an EVOS FL digital inverted fluorescence microscope (Life Technologies, #AMF4300PM) at 10X magnification in phase-contrast and GFP channels. Representative images are shown in Supplemental Fig. 2A.

EGFP quantitation was performed using a plate reader assay. HeLa cells were transfected with identical plasmids as above and plated in a 96-well clear tissue culture plate (Celltreat, #229195). n=5 transfections per condition, 2×10^4^ cells per well. Untransfected cells were plated as a negative control. After 20 hr, growth media was aspirated from the cells and replaced with 100 uL PBS. EGFP fluorescence in each well was measured on a Bio-Tek Cytation 3 plate reader (470 nm excitation, 510 nm emission, 146 gain). Baseline intensity was determined by measuring EGFP fluorescence of untransfected cells, and the mean baseline intensity was subtracted from experimental fluorescence measurements.

### Cell viability assay

The PrestoBlue resazurin assay (Invitrogen, #A13261) was used to quantify cell viability in response to CUG-targeting Cas13d. HeLa cells were transfected in a 96-well format with plasmids encoding Cas13d, CUG-targeting or non-targeting gRNA, and CUG_480_ target. n=5 transfections per condition, 2×10^4^ cells per well, with a total volume of 100 uL growth medium per well. 5 wells each of untransfected cells and media alone were also plated. 20 hr after transfection, 11 uL PrestoBlue reagent was added to each well and mixed gently by pipetting. Cells were incubated with PrestoBlue reagent at 37°C for 30 min before measuring fluorescence on a Bio-Tek Cytation 3 plate reader (550 nm excitation, 590 nm emission, 60 gain). Baseline intensity was calculated as the mean fluorescence of cell-free media and was subtracted from all measurements. Cell viability was calculated by normalizing experimental measurements by the mean fluorescence intensity of untransfected cells.

### Cas13d specificity transcriptomic analysis

RNA-seq was performed to screen for transcriptomic off-targets of repeat-targeting approaches (Cas13d, dCas9, and shRNA). CUG-targeted systems and non-targeting controls (no gRNA or non-targeting shRNA) were transfected in triplicate into HeLa cells. CUG_480_ target plasmid was omitted to enrich for off-target events. After 68 hr incubation, RNA was extracted with 300 uL TRIzol and purified using the Direct-zol RNA Miniprep Kit. Ribosomal RNA was depleted from 300 ng total RNA using the NEBNext rRNA Depletion Kit (NEB, #E6310L), and libraries were generated for next-generation sequencing using the NEBNext Ultra II Directional RNA Library Prep Kit for Illumina (NEB, #E7760L) according to the manufacturer protocol. Libraries were multiplexed with Illumina i5 and i7 barcoding primers, pooled at 4 nM, and sequenced in a 2×76 bp paired-end format on an Illumina NextSeq 500. >14 million paired end reads were sequenced per library and recorded in FASTQ format.

Transcript expression was quantified by pseudoalignment to the hg19 human reference genome using kallisto 0.43.0 (Bray et al., 2016). Reads were aligned to hg19 using HISAT2 2.0.0-beta (Kim et al., 2019). For each targeting technology (Cas13d, dCas9, shRNA), differential gene expression analysis was performed using DESeq2 1.30.1 (Love et al., 2014) to compare CUG-targeting and non-targeting conditions, with gene counts calculated by htseq-count 0.11.2 (Anders et al., 2015) provided as input. Off-targets were defined as differentially expressed (DE) genes with a false discovery rate (FDR) q<0.05. For Cas13d and shRNA libraries, gene ontology (GO) analysis was performed using PANTHER (Mi et al., 2019) (annotation version 2021-05-01) to identify biological processes associated with each set of DE genes. GO biological processes with FDR q<0.05 were considered significant and were assigned to categories according to their descriptions.

For all annotated human transcripts, longest CUG_n_ repeat tract length (in C, U, or G registers) was determined from the NCBI RefSeq reference mRNA sequence. Transcripts were grouped by maximum CUG_n_ length, and median log_2_ fold-change of TPM between CUG_n_-targeting and non-targeting conditions was calculated for each transcript group and targeting approach.

### Collateral activity mCherry fluorescence assays

A plate reader assay was developed to quantitate bulk mCherry fluorescence in transfected cells. HeLa, HEK293, or Neuro2a cells were transfected in a 96-well format with pXR001, targeting or non-targeting gRNA in pXR003, target or control plasmid, and pmCherry. n=5 transfections per condition, 2×10^4^ cells per well, with a total volume of 100 uL growth medium per well. 5 wells of untransfected cells were also plated. After 20 hr incubation, transfection media was aspirated and replaced with 100 uL PBS. mCherry fluorescence was measured on a Bio-Tek Cytation 3 plate reader (587 nm excitation, 627 nm emission, 202 gain). Baseline intensity was defined as the mean mCherry fluorescence of untransfected cells and was subtracted from experimental measurements.

For single-cell measurement of mCherry expression, HeLa cells were transfected in 4-well coated glass chamber slides with plasmids encoding Cas13d and EGFP, CUG-1 or NT gRNA, CUG_480_ target or CUG_0_ control RNA, and mCherry. After 20 hr incubation, cells were washed with PBS and fixed in 4% PFA at RT for 10 min. Cells were washed 3X with PBS, and nuclei were stained with 1 ng/uL DAPI in PBS for 5 min at RT, followed by an additional three PBS washes. Slides were mounted with glass #1.5 coverslips in Fluoroshield antifade mounting medium, sealed with clear nail polish, and stored at −20°C until imaging. Widefield epifluorescence imaging was performed on a Zeiss LSM 880 microscope with a Plan-Apochromat 40x/1.3 Oil DIC M27 objective lens. 5+ images were collected for each condition and were processed in Fiji v2.0.0-rc-69/1.52p (Schindelin et al., 2012). In each image, all EGFP-positive cells were manually segmented, and total mCherry fluorescence intensity was measured for each cell. >26 cells were measured for each condition. Distributions of mCherry expression were compared between conditions using the two-sided Mann-Whitney *U* test.

### Collateral activity RNA assay

To enable precise measurement of Cas13d collateral RNase activity when targeting transgenic or endogenous RNAs, we developed a HeLa cell line containing genomically integrated cassettes expressing mCherry constitutively and Cas13d and EGFP under a tetracycline-inducible promoter (HeLa-tet:Cas13d-mCherry). First, HeLa cells were transfected in a 12-well format (1.5×10^5^ cells) with 200 ng pB-Tet-Cas13d and 800 ng of plasmid encoding codon-optimized piggyBac transposase (mPB) (Cadiñanos and Bradley, 2007). After 2 days incubation, transfection media was replaced with 10% FBS growth media containing 2 μg/mL puromycin (AG Scientific, #P-1033-SOL) to select for transposon integration. Cells were passaged to a 10-cm dish upon reaching confluency, and puromycin selection was maintained for 2 weeks. 1.5×10^5^ cells were then passaged to a 12-well dish and transfected with 200 ng pB-mCherry and 800 ng mPB plasmid to integrate a constitutively expressing mCherry gene. Once confluent, cells were again passaged to a 10-cm dish. Two weeks after integration, Cas13d and EGFP expression was induced by incubation with 2 μM doxycycline (Sigma, #D3447-500MG) for 2 days, and EGFP-positive and mCherry-positive single cells were sorted into 96-well plates using a BD FACSAria II cell sorter at the University of Florida Interdisciplinary Center for Biotechnology Research (ICBR). Twelve EGFP+ and mCherry+ clones were manually identified using an EVOS FL microscope and expanded. The clone exhibiting the strongest expression of both EGFP and mCherry upon addition of doxycycline was chosen for the collateral activity RNA assay.

To perform the assay, HeLa-tet:Cas13d-mCherry and wild-type HeLa cells were mixed at a 1:4 ratio in growth media containing 2 μM doxycycline and plated in a 24-well plate format at a density of 7.5×10^4^ cells/well. Cells were transfected with 125 ng gRNA plasmid and either 125 ng CUG_480_ plasmid (for CUG-targeting validation experiment) or 125 ng pUC19 as inert carrier DNA (for endogenous gene targeting experiment). n=3 transfections per condition. After 44 hr, RNA was extracted with 300 uL TRIzol and purified using the Direct-zol RNA Miniprep Kit. cDNA was reverse-transcribed from 50 ng RNA using the iScript Reverse Transcription Supermix and diluted 10-fold in water. mCherry and control (*GAPDH*) expression levels were measured separately by qPCR from 4 uL of diluted cDNA using *Taq* DNA Polymerase (NEB, #M0270L), dsGreen DNA detection dye (Lumiprobe, #11010), and 200 nM each of forward and reverse primers (mCherry_F/mCherry_R and GAPDH_F/GAPDH_R, respectively). Primer sequences are presented in Supplemental Table 4. qPCR reactions were performed in duplicate on a C1000 Touch thermocycler (Bio-Rad) using a two-step cycling protocol: initial denaturation at 95°C for 3 min, followed by 40 cycles of denaturation at 95°C for 15 s and annealing and extension at 60°C for 45 s. dsGreen fluorescence was measured after the extension step of each cycle. *C_q_* values were calculated using Bio-Rad CFX Manager 3.1 software, and the difference between mCherry and *GAPDH C_q_* (Δ*C_q_*) was calculated and averaged across the two qPCR replicates. The ratio of mCherry to *GAPDH* mRNA expression (defined as 2 ^△*Cq*^) was plotted for each condition. Δ*Cq* values in targeting and non-targeting conditions were compared using the one-tailed Student’s *t* test (independent samples, equal variance).

### Modeling of negative autoregulation by gRNA excision

An ordinary differential equation (ODE) model was constructed that describes the dynamics of GENO autoregulation of Cas13d expression:

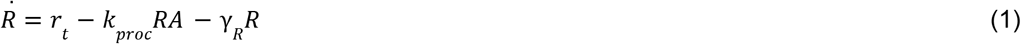

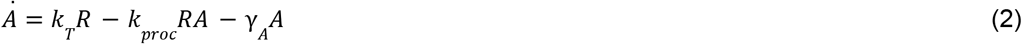

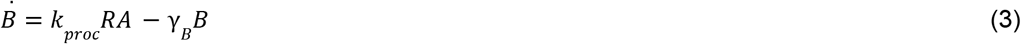

where *R* is the concentration of Cas13d mRNA (also containing the pre-crRNA), *A* is the concentration of Cas13d apoprotein, *B* is the concentration of Cas13d:gRNA binary complex, *r_t_* is the rate of RNA polymerase II transcription of the Cas13d mRNA, *k_T_* is the translation rate constant of Cas13d, *k_proc_* is the rate constant of crRNA processing, and *γ_i_* is the degradation rate constant of species *i.* Transcription is modeled by zero-order kinetics, translation and degradation by first-order kinetics, and crRNA processing by second-order kinetics. Further details of the dynamical model, a reference model for unregulated Cas13d, and analytical proofs are presented in Supplemental Note.

To calculate equilibrium binary complex concentration across ranges of biochemical parameters, simulations were conducted by ODE integration using Python 3. RNA and protein degradation rates were estimated from median half-lives determined from the literature (10 hr (Yang et al., 2003) and 36 hr (Cambridge et al., 2011), respectively), and the degradation rates of Cas13d apoprotein and binary complex were estimated to be equivalent. In the unregulated reference model, the half-life of free gRNA was estimated as 2 hr. For each vector of parameters *r_t_, k_T_,* and *k_proc_,* the dynamical model was integrated over 480 hr with 5000 timesteps to reach steady state. Autoregulation efficiency *η*_GENO_ was calculated for each parameter vector as the difference of unity and the ratio of equilibrium binary complex concentrations between GENO and unregulated conditions (see Supplemental Note).

### Western blot

Cas13d protein expression in unregulated and autoregulated conditions was measured by Western blot. HeLa cells were transfected in 12-well format with 500 ng Cas13d plasmid (unregulated or GENO, +100 ng NT gRNA plasmid for unregulated Cas13d). n=3 transfections per condition. After 44 hr incubation, cells were washed with 1 mL PBS, and protein was extracted for 15 min on ice in 150 uL radioimmunoprecipitation assay (RIPA) buffer (Thermo Scientific, #89901) containing 2X SIGMAFAST Protease Inhibitor Cocktail (Sigma-Aldrich, #S8830) and 1 mM phenylmethylsulfonyl fluoride (Sigma-Aldrich, #10837091001). Extractions were transferred to microcentrifuge tubes and centrifuged at 12,000 RCF for 15 min at 4°C. Total protein content in supernatant was measured using the Pierce BCA Protein Assay (Thermo Scientific, #23225). For each sample, 10 μg protein was mixed with 5.4 uL 4X NuPage LDS Sample Buffer (Invitrogen, #NP0008), 0.6 uL 100 mM dithiothreitol (NEB, #B1034A), and water to a total volume of 24 uL. Protein was denatured at 95°C for 5 min prior to loading on a NuPage 4-12% Bis-Tris polyacrylamide gel (#NP0336), with 3 uL Precision Plus Protein All Blue prestained standards (Bio-Rad, #1610373) loaded as a ladder. Gel electrophoresis was performed at 120 V for 45 min in MOPS running buffer (10.46 mg/mL 3-(N-morpholino)propanesulfonic acid (MOPS), 6.06 mg/mL Tris base, 1 mg/mL SDS, 0.3 mg/mL ethylenediaminetetraacetic acid (EDTA) in water). Protein was transferred to a methanol-activated polyvinylidene fluoride (PVDF) membrane (Bio-Rad, #1620264) using the iBlot 2 dry transfer system (ThermoFisher Scientific, #IB21001).

The membrane was incubated in 5 mL SEA BLOCK Blocking Buffer (Thermo Scientific, #37527) for 30 min at RT in a dark container on a rocking mixer. Blocking buffer was removed and replaced with primary antibody solution (rabbit α-HA (C29F4, 1:1000, Cell Signaling Technology) and mouse α-HSP70 (5A5, 1:500, Invitrogen) in 5 mL SEA BLOCK + 0.2% Tween 20 (Fisher Bioreagents, #BP152-1)), and the membrane was placed in a 4°C room on a rocking mixer overnight. The next day, the primary antibody solution was removed, and the membrane was washed 3X in PBS-T. After washing, the secondary antibody solution was added (IRDye 800CW donkey α-rabbit and 680RD donkey α-mouse (1:5000 each, LI-COR) in 5 mL SEA BLOCK + 0.2% Tween 20) and incubated for 1 hr at RT on a rocking mixer. The secondary antibodies were removed, and the membrane was washed 3X in PBS-T. The blot was imaged on a LI-COR Odyssey CLx scanner at 700 and 800 nm excitation wavelengths. Image processing was performed in Fiji, and blot images were inverted for publication.

### AAV treatment of DM1 myoblasts

Recombinant AAV constructs were packaged in AAV6 capsids and purified by SignaGen Laboratories (#SL100865, >10^13^ VG/mL). Production of AAV-CMV-unreg-CUG1 was attempted on three occasions but titers >10^9^ VG/mL could not be obtained. DM1 myoblasts were grown to 50% confluency in 4-well coated glass chamber slides and transduced with AAV at a titer of 2×10^10^ VG/mL for 6 days prior to fixation and staining.

### Hybridization chain reaction FISH (HCR FISH)

To image single molecules of Cas13d mRNA by FISH, HCR v3.0 was performed for signal amplification (Choi et al., 2018). A custom HCR probe set matching the Cas13d CDS mRNA sequence (30 split-initiator probes, B1 amplifier) was designed by Molecular Instruments (Los Angeles, CA). After AAV transduction and culture, myoblasts were fixed in 4% PFA for 10 min at RT, washed 3X with PBS, permeabilized with 0.2% Triton X-100 in PBS for 10 min at RT, and washed again 3X with PBS. If IF was also performed, cells were incubated with IF blocking buffer for 30 min prior to incubation of primary antibody (rabbit α-HA (C29F4, 1:1000)) in IF blocking buffer with cells at 4°C overnight. The next day, cells were washed 3X with IF wash buffer and incubated with secondary antibody (α-rabbit Alexa Fluor 568, 1:1000) in IF blocking buffer for 1 hr at RT, followed by three washes in PBS. To cross-link antibodies to antigens prior to HCR FISH, cells were fixed again in 4% PFA for 10 min and washed 3X with PBS.

For HCR FISH, cells were washed twice with 2X SSC and incubated with HCR Probe Hybridization Buffer (Molecular Instruments) for 30 min at 37°C. For each well, 1.2 pmol of Cas13d HCR probe set was mixed with 300 uL HCR Probe Hybridization Buffer and incubated with cells overnight at 37°C in a humidified chamber. If CUG_n_ FISH was also performed, CAG_10_ Alexa Fluor 594 probe was mixed with the HCR probe solution at 380 ng/mL and prior to overnight incubation. The following day, cells were washed 4X with HCR Probe Wash Buffer (Molecular Instruments) at 37°C and twice with 5X SSC + 0.1% Tween 20 (5X SSCT) at RT to remove excess probe, and cells were equilibrated with HCR Amplification Buffer (Molecular Instruments) for 30 min at RT. To prepare amplification hairpins, 2.4 pmol each of B1 h1 and B1 h2 hairpins (Molecular Instruments, 647 nm) were heated in separate tubes to 95°C for 90 s and cooled to RT in a light-protected compartment for 30 min. Hairpins were mixed in 40 uL HCR Amplification Buffer and applied to cells at RT for 2 hr. Cells were washed 5 times in 5X SSCT and 3 times in PBS to remove excess hairpins. For nucleus and cytoplasm staining, 1 ng/uL DAPI and 0.1X HCS CellMask Green (ThermoFisher Scientific, #H32714) in PBS was added to the cells for 5 min at RT, followed by three washes with PBS. Slides were mounted on glass #1.5 coverslips using ProLong Diamond antifade mounting medium (ThermoFisher Scientific, #P36965) and stored at −20°C until imaging. 2D confocal imaging was performed on a Zeiss LSM 880 microscope with Plan-Apochromat 40x/1.3 Oil DIC M27 and Plan-Apochromat 20x/0.8 M27 objective lenses. All fields of view were chosen randomly in the DAPI channel.

For cells labeled with Cas13d HCR FISH and α-HA IF, 3-5 images per condition were collected at 20x magnification. All images were processed in Python 3 to analyze HCR FISH and IF signal intensities per nucleus. Images in CZI format were loaded and channels were separated into 2D arrays. Nuclei were identified by thresholding the DAPI channel using a modified Otsu’s method (Otsu, 1979). Within each nucleus, the mean pixel intensity in each of the HCR FISH and IF channels was calculated. Nuclear measurements were pooled across all images within each condition.

For cells labeled with Cas13d HCR FISH and CUG_n_ FISH, 21 images per condition were collected at 40x magnification and processed using Python 3 to measure FISH intensities and detect diffraction-limited spots. Images were loaded and nuclei were thresholded as above. Mean nuclear HCR FISH and CUG_n_ FISH signal intensities were calculated for each nucleus. In both FISH channels, spots were detected using the Laplacian of Gaussian method (Lindeberg, 1998) (thresholds of 0.6% for HCR FISH and 0.2% for CUG_n_ FISH, Gaussian standard deviation of 2 px). For each detected CUG_n_ FISH spot, spot intensity was calculated by convolving a Gaussian kernel (standard deviation of 2 px) with the FISH channel image and measuring the pixel brightness at the centroid of the spot in the blurred image. Per-nucleus and per-spot measurements were pooled across all images within each condition.

## Code availability

All code developed for this work was written in Python and is publicly available on GitHub at https://github.com/cpkelley94/geno.

## Data availability

RNA-seq raw read data (FASTQ format) and kallisto transcript expression table are available through the Gene Expression Omnibus (GEO) at accession no. GSE191329. Raw microscopy images (CZI format) are available upon request.

