## Supplemental Note for "Negative autoregulation mitigates collateral RNase activity of repeat-targeting CRISPR-Cas13d in mammalian cells"

### Supplemental Note: Dynamical model of Cas13d gRNA excision for negative-autoregulatory optimization

University of Florida

November 19, 2021

#### 1 Model description

The following model is proposed to describe the kinetics and equilibria of a Cas13d negative autoregulation strategy mediated by gRNA excision (GENO):

$$\dot{R} = \text{transcription} - \text{processing} - \text{degradation}$$
$$\dot{A} = \text{translation} - \text{processing} - \text{degradation}$$
$$\dot{B} = \text{processing} - \text{degradation}$$

This description relies on the following assumptions:

1. A deterministic model describes the mean dynamics of the underlying stochastic process with reasonable accuracy.
2. gRNA processing is performed by apoprotein only, upon Cas13d:gRNA binary complex formation.
3. Nascent Cas13d translation products quickly diffuse away from the domain of their mRNAs.
4. Cas13d:gRNA binary complex formation is irreversible.
5. Nuclear/cytoplasmic compartmentalization affects crRNA processing negligibly.

These biochemical dynamics and assumptions produce the following differential equation model (GENO):

$$\dot{R} = r_t - k_p RA - \gamma_R R \tag{1.1}$$
$$\dot{A} = k_T R - k_p RA - \gamma_A A \tag{1.2}$$
$$\dot{B} = k_p RA - \gamma_B B \tag{1.3}$$

where:

- $R$: concentration of Cas13d mRNA
- $A$: concentration of Cas13d apoprotein
- $B$: concentration of Cas13d:gRNA binary complex
- $r_t$: rate of Pol II transcription of Cas13d mRNA
- $k_T$: rate of translation of Cas13d protein
- $k_p$: rate of crRNA processing
- $\gamma_i$: rate of degradation of species $i$

A reference model (REF) is also presented for comparison, in which autoregulation is absent and gRNA processing and Cas13d expression are independent:

$$\dot{R} = r_t - \gamma_R R \tag{1.4}$$
$$\dot{A} = k_T R - k_p G A - \gamma_A A \tag{1.5}$$
$$\dot{B} = k_p G A - \gamma_B B \tag{1.6}$$
$$\dot{G} = r_G - k_p G A - \gamma_G G \tag{1.7}$$

where:

- $G$: concentration of unbound gRNA
- $r_G$: rate of Pol III transcription of gRNA

### 2 Equilibrium analysis

#### 2.1 GENO model

At equilibrium, the GENO model reduces to the following:

$$0 = r_t - k_p \hat{R}\hat{A} - \gamma_R \hat{R} \tag{2.1}$$
$$0 = k_T \hat{R} - k_p \hat{R}\hat{A} - \gamma_A \hat{A} \tag{2.2}$$
$$0 = k_p \hat{R}\hat{A} - \gamma_B \hat{B} \tag{2.3}$$

From (2.1),

$$\hat{A} = \frac{r_t - \gamma_R \hat{R}}{k_p \hat{R}} \tag{2.4}$$

By substitution of (2.4) into (2.2),

$$\frac{r_t - \gamma_R \hat{R}}{k_p \hat{R}} = \frac{k_T \hat{R}}{k_p \hat{R} + \gamma_A} \tag{2.5}$$

which reduces to the quadratic equation

$$0 = k_p(k_T + \gamma_R)\hat{R}^2 - (k_p r_t - \gamma_A \gamma_R)\hat{R} - \gamma_A r_t \tag{2.6}$$

The solutions of this quadratic equation are

$$\hat{R} = \frac{k_p r_t - \gamma_A \gamma_R \pm \sqrt{(k_p r_t - \gamma_A \gamma_R)^2 + 4\gamma_A r_t k_p (k_T + \gamma_R)}}{2k_p(k_T + \gamma_R)} \tag{2.7}$$

By inspection,

$$\left| k_p r_t - \gamma_A \gamma_R \right| < \sqrt{(k_p r_t - \gamma_A \gamma_R)^2 + 4\gamma_A r_t k_p (k_T + \gamma_R)} \tag{2.8}$$

Therefore, regardless of the sign of the term $\left| k_p r_t - \gamma_A \gamma_R \right|$, the quadratic equation (2.6) has a single positive solution given by

$$\hat{R} = \frac{k_p r_t - \gamma_A \gamma_R + \sqrt{(k_p r_t - \gamma_A \gamma_R)^2 + 4\gamma_A r_t k_p (k_T + \gamma_R)}}{2k_p(k_T + \gamma_R)} \tag{2.9}$$

Equivalently,

$$\hat{R} = \frac{r_t}{\gamma_R} \left( \frac{k_p \gamma_R - \frac{\gamma_A \gamma_R^2}{r_t} + \sqrt{\left( k_p \gamma_R - \frac{\gamma_A \gamma_R^2}{r_t} \right)^2 + \frac{4k_p k_T \gamma_A \gamma_R^2}{r_t}}}{2k_p(k_T + \gamma_R)} \right) \tag{2.10}$$

From (2.3) and (2.4),

$$\hat{B} = \frac{r_t - \gamma_R \hat{R}}{\gamma_B} \tag{2.11}$$

Thus, the system described by the GENO model has a single equilibrium point ($\hat{R}^{\text{GENO}}$, $\hat{A}^{\text{GENO}}$, $\hat{B}^{\text{GENO}}$) at the solutions provided in (2.10), (2.4), and (2.11).

#### 2.2 REF model

At equilibrium, the REF model reduces to the following:

$$0 = r_t - \gamma_R \hat{R} \tag{2.12}$$
$$0 = k_T \hat{R} - k_p \hat{G}\hat{A} - \gamma_A \hat{A} \tag{2.13}$$
$$0 = k_p \hat{G}\hat{A} - \gamma_B \hat{B} \tag{2.14}$$
$$0 = r_G - k_p \hat{G}\hat{A} - \gamma_G \hat{G} \tag{2.15}$$

From (2.12),

$$\hat{R} = \frac{r_t}{\gamma_R} \tag{2.16}$$

From (2.13) and (2.16),

$$\hat{A} = \frac{k_T r_t}{\gamma_R}\left(\frac{1}{k_p \hat{G} + \gamma_A}\right) \tag{2.17}$$

From (2.15),

$$\hat{G} = \frac{r_G}{k_p \hat{A} + \gamma_G} \tag{2.18}$$

Substituting (2.18) into (2.17) yields

$$\hat{A} = \frac{k_T r_t}{\gamma_R}\left(\frac{1}{\frac{k_p r_G}{k_p \hat{A} + \gamma_G} + \gamma_A}\right) \tag{2.19}$$

which simplifies to the quadratic equation

$$\gamma_A k_p \hat{A}^2 + \left(k_p r_G + \gamma_A \gamma_G - \frac{k_T r_t k_p}{\gamma_R}\right)\hat{A} - \frac{k_T r_t \gamma_G}{\gamma_R} = 0 \tag{2.20}$$

As the product of the first and third coefficients is strictly negative, the discriminant is strictly positive. Thus, the polynomial has a single positive real solution for $\hat{A}$.

From (2.14),

$$\hat{B} = \frac{k_p \hat{G} \hat{A}}{\gamma_B} \tag{2.21}$$

Substituting (2.17) into (2.21) yields

$$\hat{B} = \frac{k_T r_t}{\gamma_R \gamma_B}\left(\frac{k_p \hat{G}}{k_p \hat{G} + \gamma_A}\right) \tag{2.22}$$

Thus, the system described by the REF model has a single equilibrium point ($\hat{R}^{\text{REF}}, \hat{A}^{\text{REF}}, \hat{B}^{\text{REF}}, \hat{G}^{\text{REF}}$) with the solution fully constrained by (2.16), (2.20), (2.22), and (2.18), respectively.

Importantly, in the REF model, the concentration of binary complex $\hat{B}$ takes the form of a Hill function, where $\lim_{\hat{G}\to 0} \hat{B} = 0$ and $\lim_{\hat{G}\to\infty} \hat{B} = \frac{k_T r_t}{\gamma_R \gamma_B}$.

The autoregulation efficiency $\eta_{\text{GENO}}$ is defined as

$$\eta_{\text{GENO}} = \frac{\hat{B}^{\text{REF}} - \hat{B}^{\text{GENO}}}{\hat{B}^{\text{REF}}} \tag{2.23}$$

### 3 Proofs

**Theorem 1.** At equilibrium, negative autoregulation by gRNA excision reduces the expression of the Cas13d mRNA compared to the reference model.

*Proof.* Proof by contradiction. The equilibrium mRNA concentration in the GENO model is provided in (2.10):

$$\hat{R}^{\text{GENO}} = \frac{r_t}{\gamma_R}\left(\frac{k_p\gamma_R - \frac{\gamma_A\gamma_R^2}{r_t} + \sqrt{\left(k_p\gamma_R - \frac{\gamma_A\gamma_R^2}{r_t}\right)^2 + \frac{4k_pk_T\gamma_A\gamma_R^2}{r_t}}}{2k_p(k_T+\gamma_R)}\right)$$

In the REF model, the equilibrium concentration is provided in (2.16):

$$\hat{R}^{\text{REF}} = \frac{r_t}{\gamma_R}$$

Assume $\hat{R}^{\text{GENO}} \geq \hat{R}^{\text{REF}}$. Equivalently,

$$\frac{k_p\gamma_R - \frac{\gamma_A\gamma_R^2}{r_t} + \sqrt{\left(k_p\gamma_R - \frac{\gamma_A\gamma_R^2}{r_t}\right)^2 + \frac{4k_pk_T\gamma_A\gamma_R^2}{r_t}}}{2k_p(k_T+\gamma_R)} \geq 1$$

$$k_p\gamma_R - \frac{\gamma_A\gamma_R^2}{r_t} + \sqrt{\left(k_p\gamma_R - \frac{\gamma_A\gamma_R^2}{r_t}\right)^2 + \frac{4k_pk_T\gamma_A\gamma_R^2}{r_t}} \geq 2k_pk_T + 2k_p\gamma_R$$

$$\sqrt{\left(k_p\gamma_R - \frac{\gamma_A\gamma_R^2}{r_t}\right)^2 + \frac{4k_pk_T\gamma_A\gamma_R^2}{r_t}} \geq 2k_pk_T + k_p\gamma_R + \frac{\gamma_A\gamma_R^2}{r_t}$$

$$\left(k_p\gamma_R - \frac{\gamma_A\gamma_R^2}{r_t}\right)^2 + \frac{4k_pk_T\gamma_A\gamma_R^2}{r_t} \geq \left(2k_pk_T + k_p\gamma_R + \frac{\gamma_A\gamma_R^2}{r_t}\right)^2$$

$$\left(\frac{\gamma_A\gamma_R^2}{r_t}\right)^2 + \frac{2\gamma_A\gamma_R^3k_p}{r_t} + k_p^2\gamma_R^2 + \frac{4k_pk_T\gamma_A\gamma_R^2}{r_t} \geq \left(\frac{\gamma_A\gamma_R^2}{r_t}\right)^2 + \frac{2\gamma_A\gamma_R^3k_p}{r_t} + k_p^2\gamma_R^2 + \frac{4k_pk_T\gamma_A\gamma_R^2}{r_t} + 4k_p^2k_T\gamma_R + 4k_p^2k_T^2$$

$$0 \geq 4k_p^2k_T\gamma_R + 4k_p^2k_T^2$$

$$0 \geq 4k_p^2k_T(\gamma_R + k_T)$$

This inequality cannot be satisfied, as all biological parameters in the right-hand term are strictly positive. Therefore, $\hat{R}^{\text{GENO}} < \hat{R}^{\text{REF}}$.

□

**Theorem 2.** Assume that gRNA is highly expressed in the reference model and is present in excess ($\hat{G} >> \gamma_A/k_p$), and that Cas13d protein translation is faster than mRNA degradation ($k_T > \gamma_R$). At equilibrium, negative autoregulation by gRNA excision reduces the concentration of active Cas13d:gRNA binary complex compared to the reference model.

*Proof.* Proof by contradiction. The equilibrium binary complex concentration in the GENO model is provided in (2.11):

$$\hat{B}^{\text{GENO}} = \frac{r_t - \gamma_R \hat{R}^{\text{GENO}}}{\gamma_B}$$

In the REF model, the equilibrium concentration is provided in (2.22):

$$\hat{B}^{\text{REF}} = \frac{k_T r_t}{\gamma_R \gamma_B} \left( \frac{k_p \hat{G}^{\text{REF}}}{k_p \hat{G}^{\text{REF}} + \gamma_A} \right)$$

If gRNA is expressed in excess in the REF model, the binary complex equilibrium concentration approaches its maximum value:

$$\lim_{\hat{G} \to \infty} \hat{B}^{\text{REF}} = \frac{k_T r_t}{\gamma_R \gamma_B}$$

Assume $\hat{B}^{\text{GENO}} \geq \lim_{\hat{G} \to \infty} \hat{B}^{\text{REF}}$. Equivalently,

$$\begin{aligned} \frac{r_t - \gamma_R \hat{R}^{\text{GENO}}}{\gamma_B} &\geq \frac{k_T r_t}{\gamma_R \gamma_B} \\ r_t - \gamma_R \hat{R}^{\text{GENO}} &\geq \frac{k_T r_t}{\gamma_R} \\ 1 - \frac{k_T}{\gamma_R} &\geq \frac{\gamma_R \hat{R}^{\text{GENO}}}{r_t} \end{aligned}$$

The right-hand term of this inequality is strictly positive. However, as $k_T > \gamma_R$, the left-hand term is strictly negative. This inequality cannot be satisfied. Thus, $\hat{B}^{\text{GENO}} < \lim_{\hat{G} \to \infty} \hat{B}^{\text{REF}}$, under the following conditions:

1. gRNA expression in the reference system is high and in excess.

2. On average, more than one Cas13d protein molecule is translated from each Cas13d mRNA in the reference system.

□
